## Supplementary Figures and Supplementary methods for "Make-or-break prime editing for bacterial genome engineering"

### Supplementary Information

#### Supplementary Figures

##### Supplementary Figure 1.

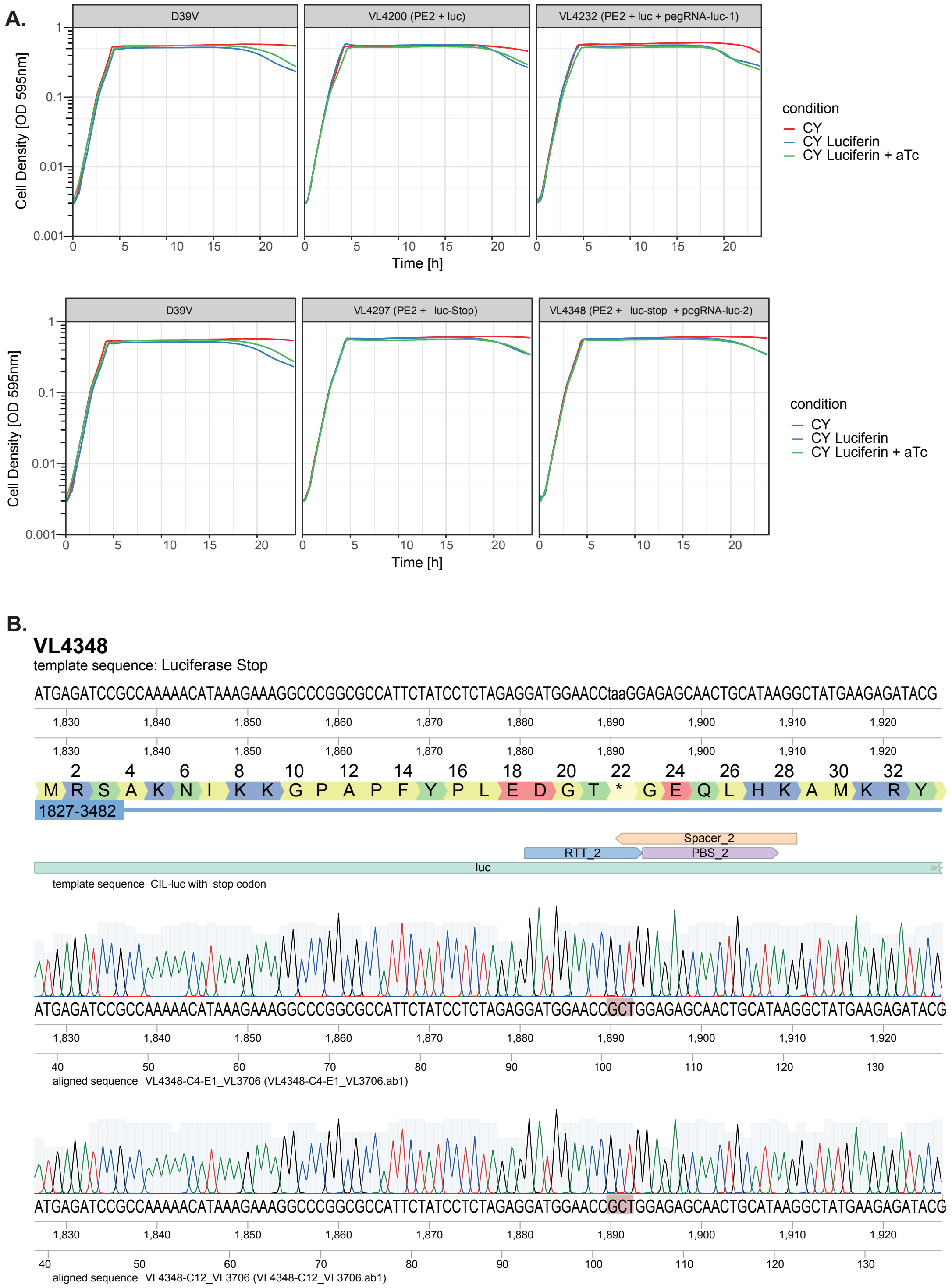

**B.**

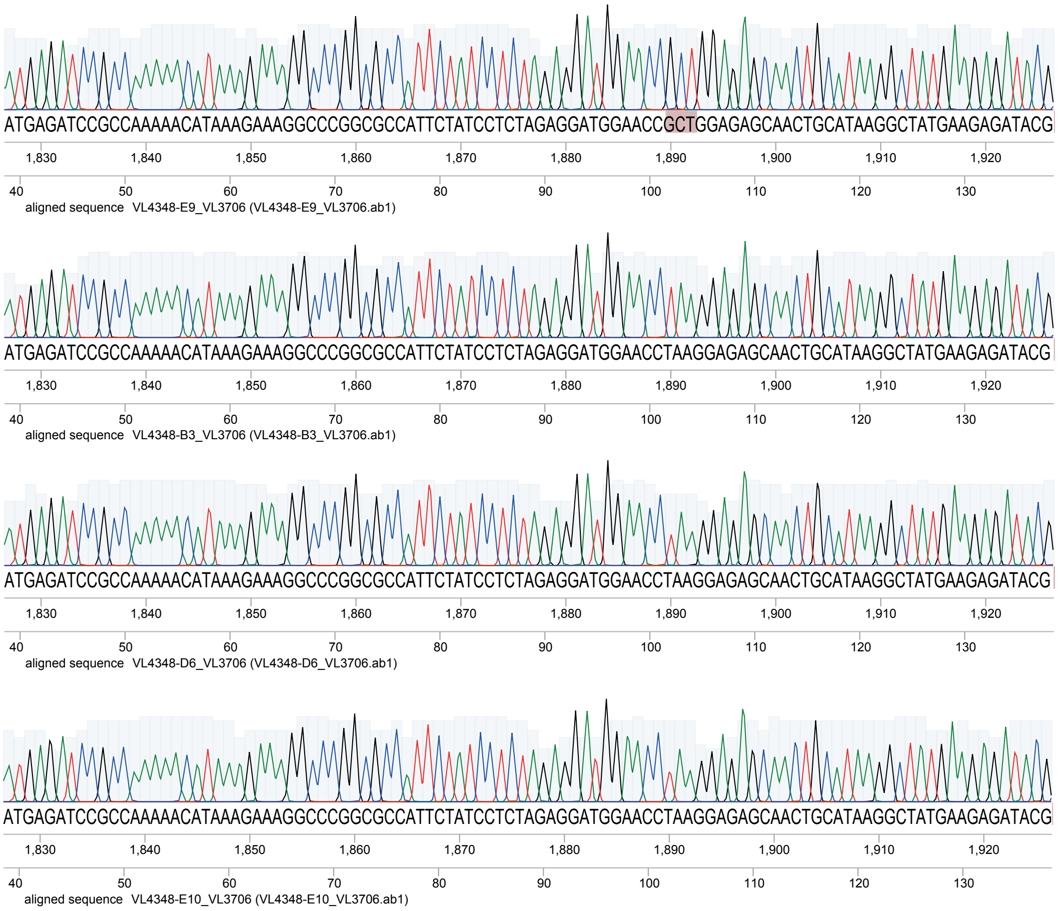
ATGAGATCCGCCAAAAACATAAAGAAAGGCCCGGCGCCATTCTATCCTCTAGAGGATGGAACCGCTGGAGAGCAACTGCATAAGGCTATGAAGAGATACG
1,830 1,840 1,850 1,860 1,870 1,880 1,890 1,900 1,910 1,920
40 50 60 70 80 90 100 110 120 130
aligned sequence VL4348-E9_VL3706 (VL4348-E9_VL3706.ab1)
ATGAGATCCGCCAAAAACATAAAGAAAGGCCCGGCGCCATTCTATCCTCTAGAGGATGGAACCTAAGGAGAGCAACTGCATAAGGCTATGAAGAGATACG
1,830 1,840 1,850 1,860 1,870 1,880 1,890 1,900 1,910 1,920
40 50 60 70 80 90 100 110 120 130
aligned sequence VL4348-B3_VL3706 (VL4348-B3_VL3706.ab1)
ATGAGATCCGCCAAAAACATAAAGAAAGGCCCGGCGCCATTCTATCCTCTAGAGGATGGAACCTAAGGAGAGCAACTGCATAAGGCTATGAAGAGATACG
1,830 1,840 1,850 1,860 1,870 1,880 1,890 1,900 1,910 1,920
40 50 60 70 80 90 100 110 120 130
aligned sequence VL4348-D6_VL3706 (VL4348-D6_VL3706.ab1)
ATGAGATCCGCCAAAAACATAAAGAAAGGCCCGGCGCCATTCTATCCTCTAGAGGATGGAACCTAAGGAGAGCAACTGCATAAGGCTATGAAGAGATACG
1,830 1,840 1,850 1,860 1,870 1,880 1,890 1,900 1,910 1,920
40 50 60 70 80 90 100 110 120 130
aligned sequence VL4348-E10_VL3706 (VL4348-E10_VL3706.ab1)

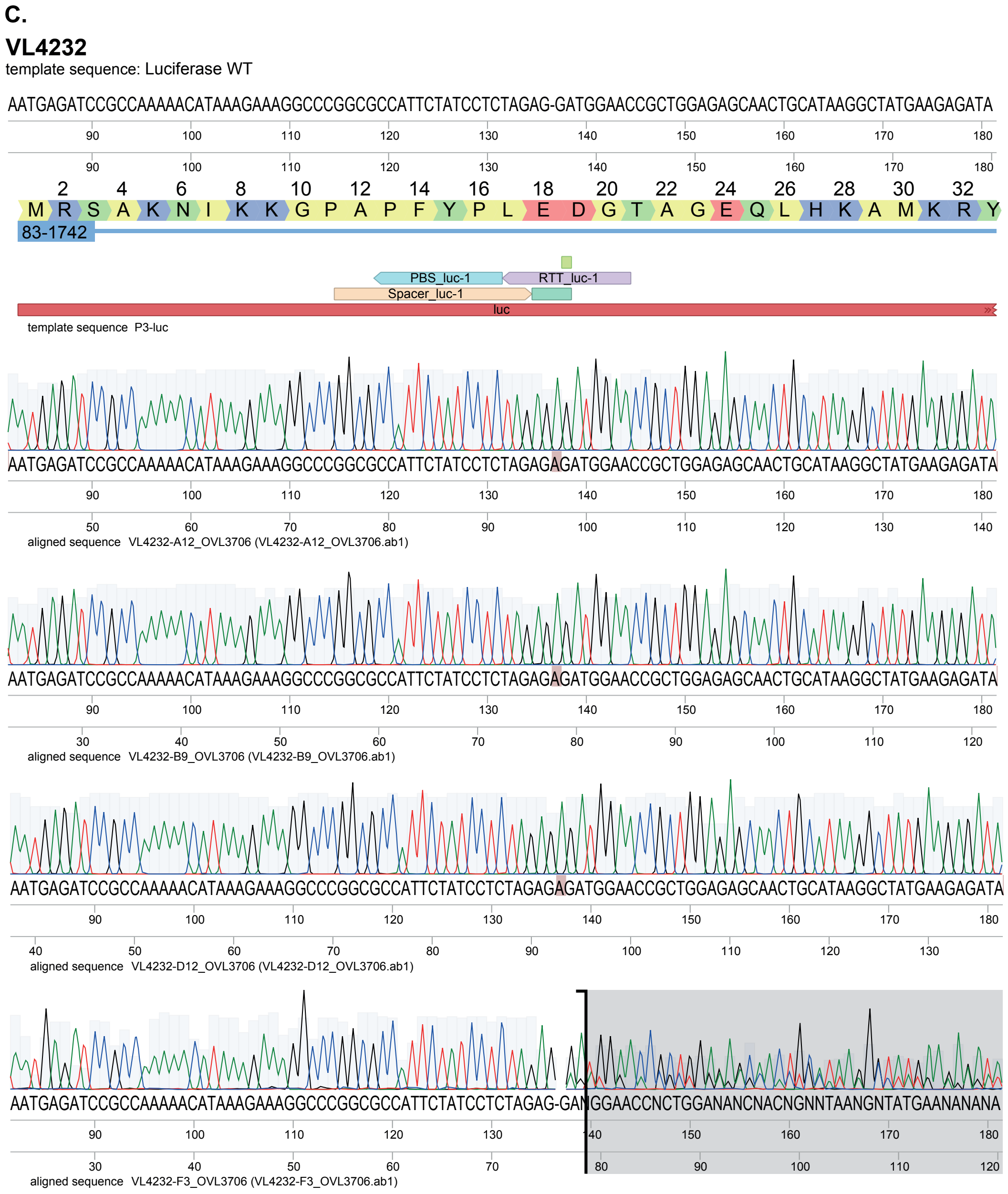


**Supplementary Figure S1.** (**A**) Induction of PE2$^{S.pn}$ together with targeting pegRNAs does not result in significant growth delay. (**B-C**) Sanger sequencing shows that non-bioluminescent clones are unedited while bioluminescent clones are edited at the targeted site for strain VL4348 (B) and vice versa for strain VL4232 (C). Note that we regularly observed mixtures of edited and non-edited clones, and these were scored as unedited.

#### Supplementary Figure 2.

A.

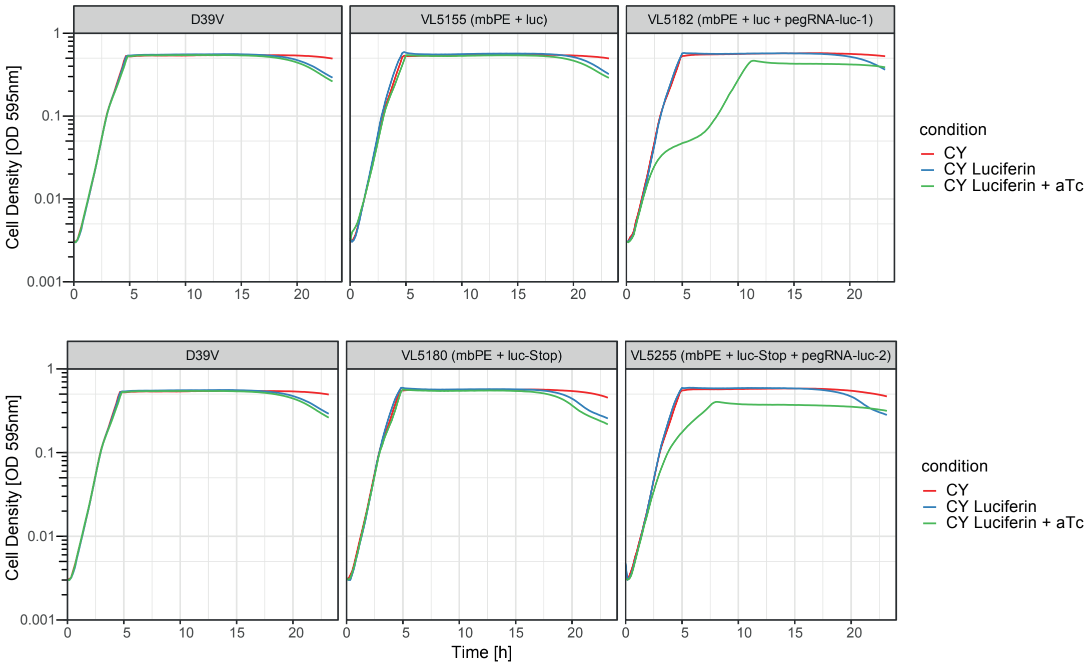


B.

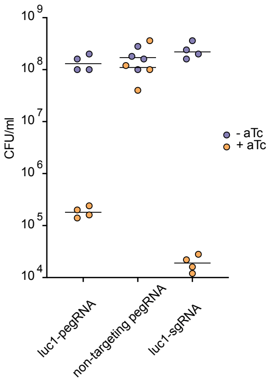

## C. VL5182

template sequence: Luciferase wild-type

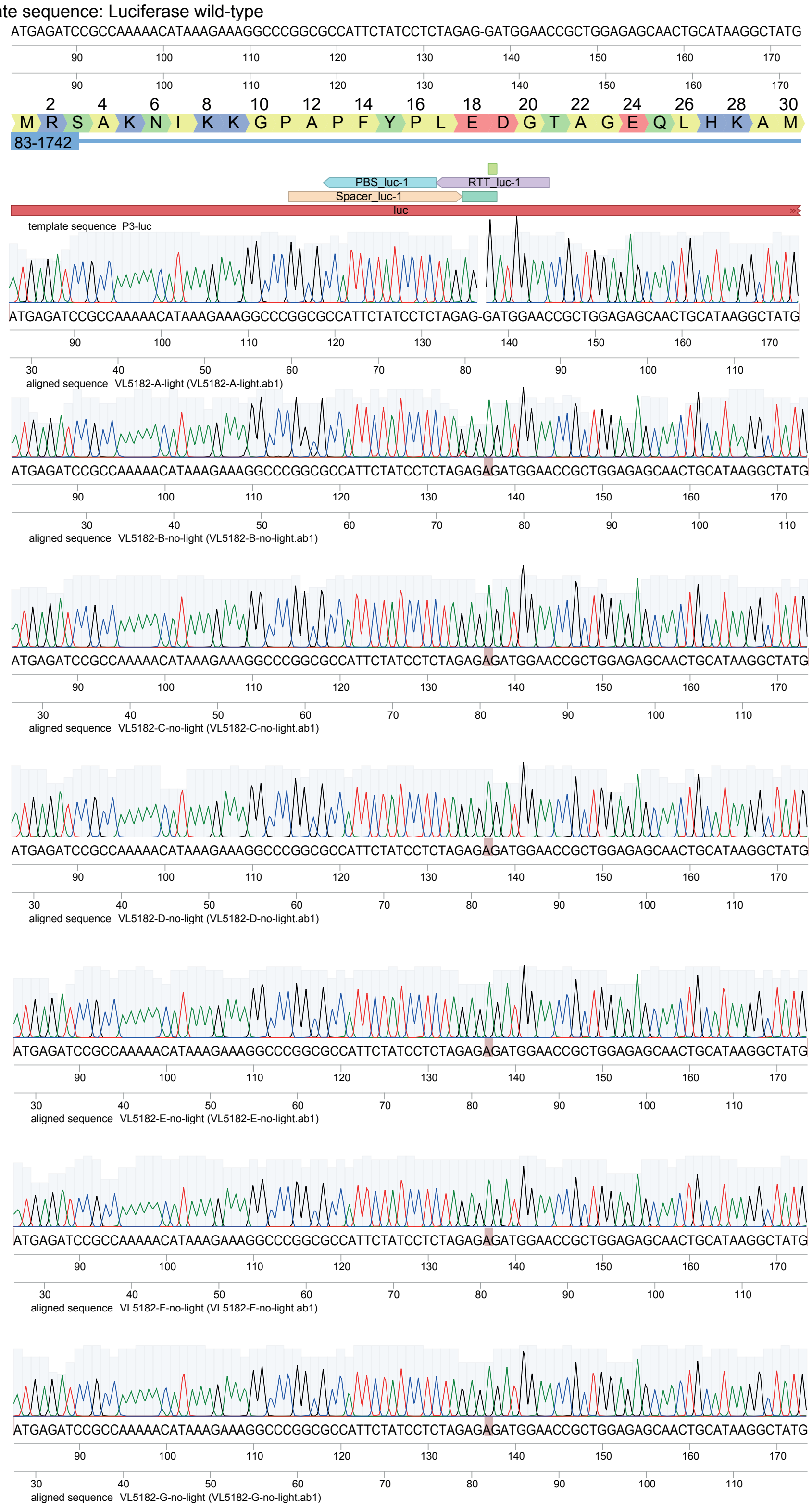

## C. VL5182

template sequence: Luciferase wild-type

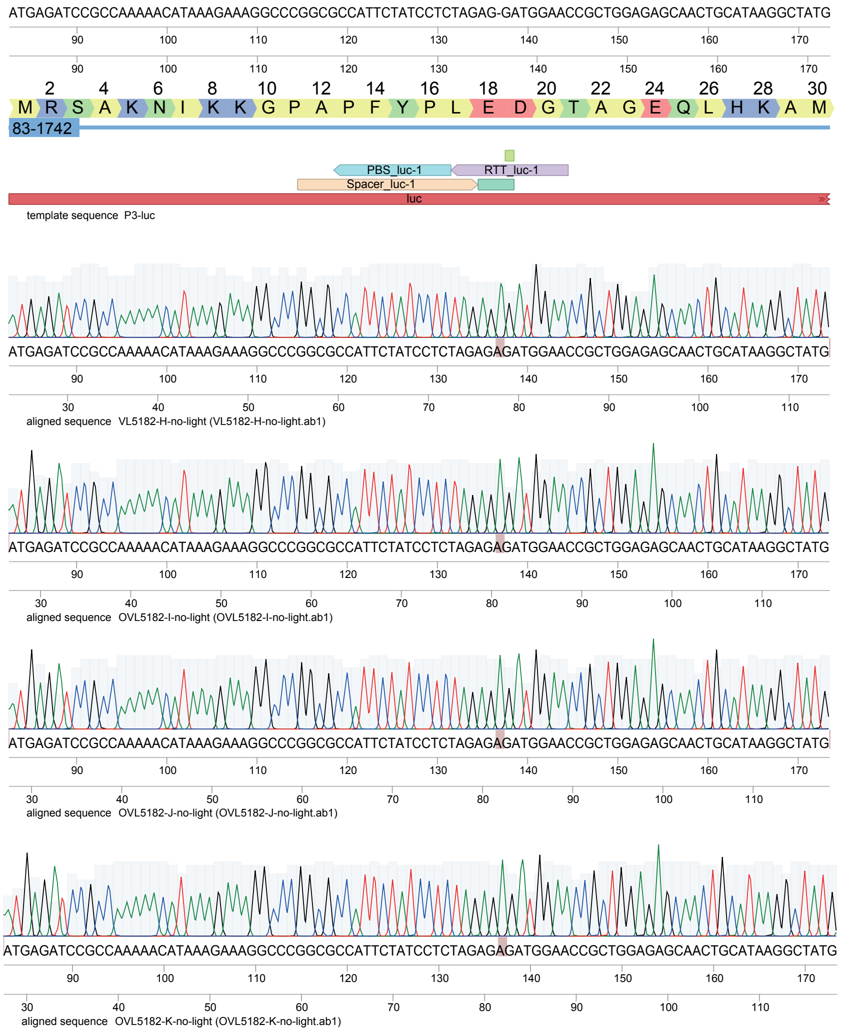

**D. VL5181**

template sequence: Luciferase wild-type

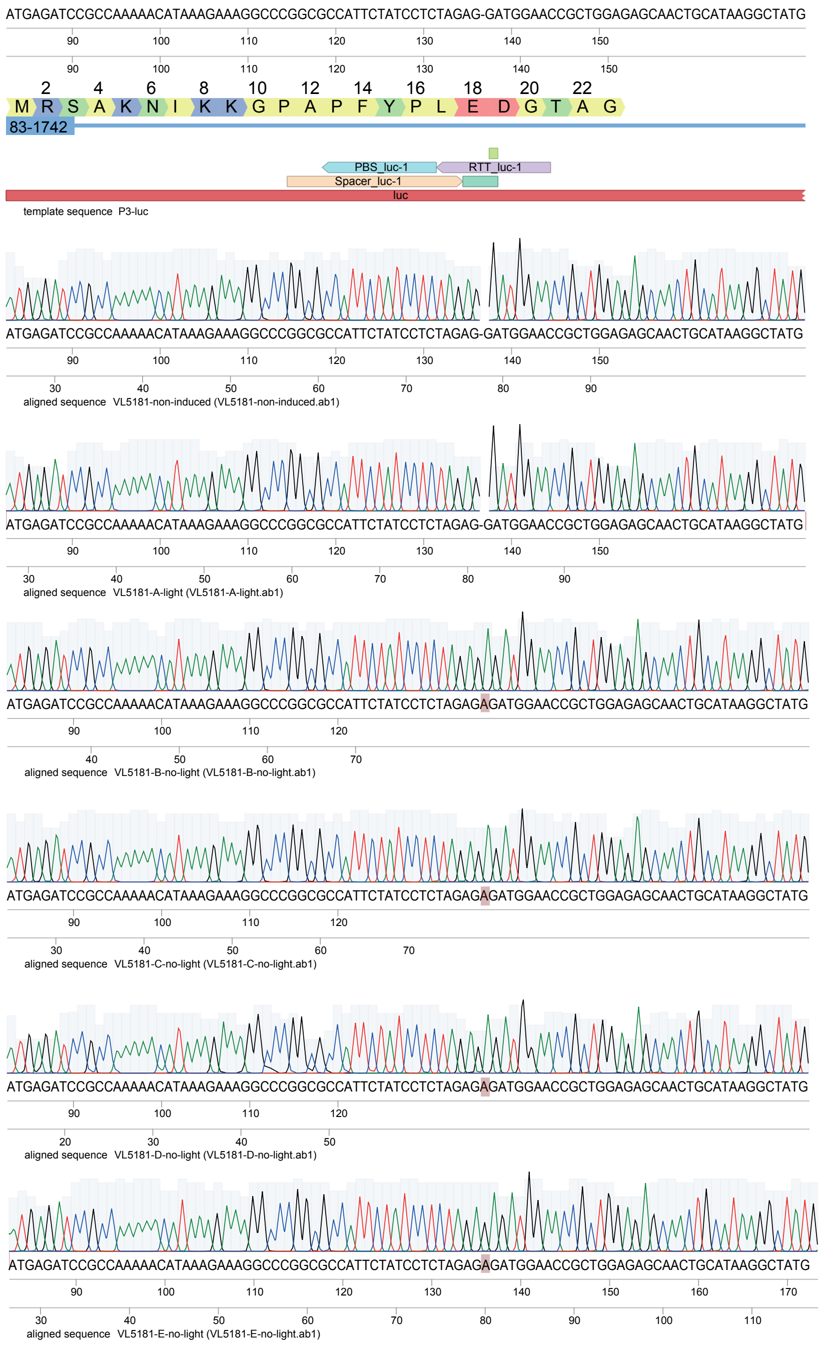

## D. VL5181

template sequence: Luciferase wild-type

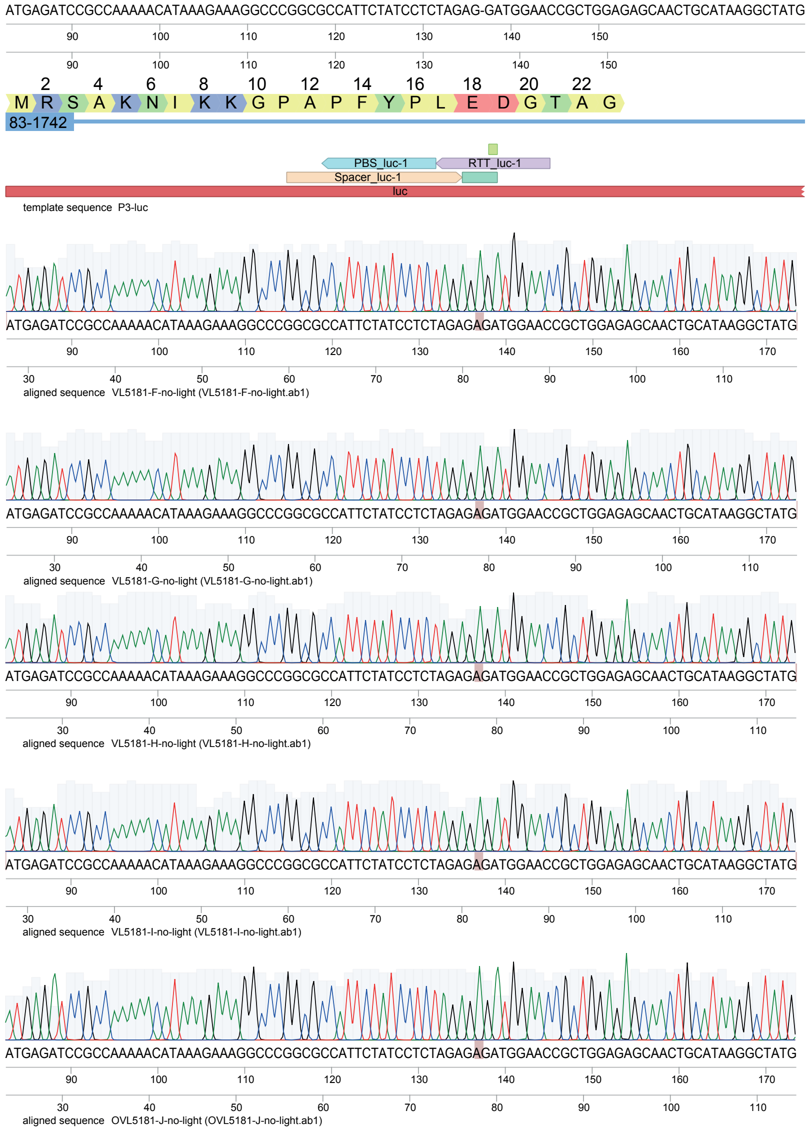

# E. VL5254

template sequence: Luciferase STOP

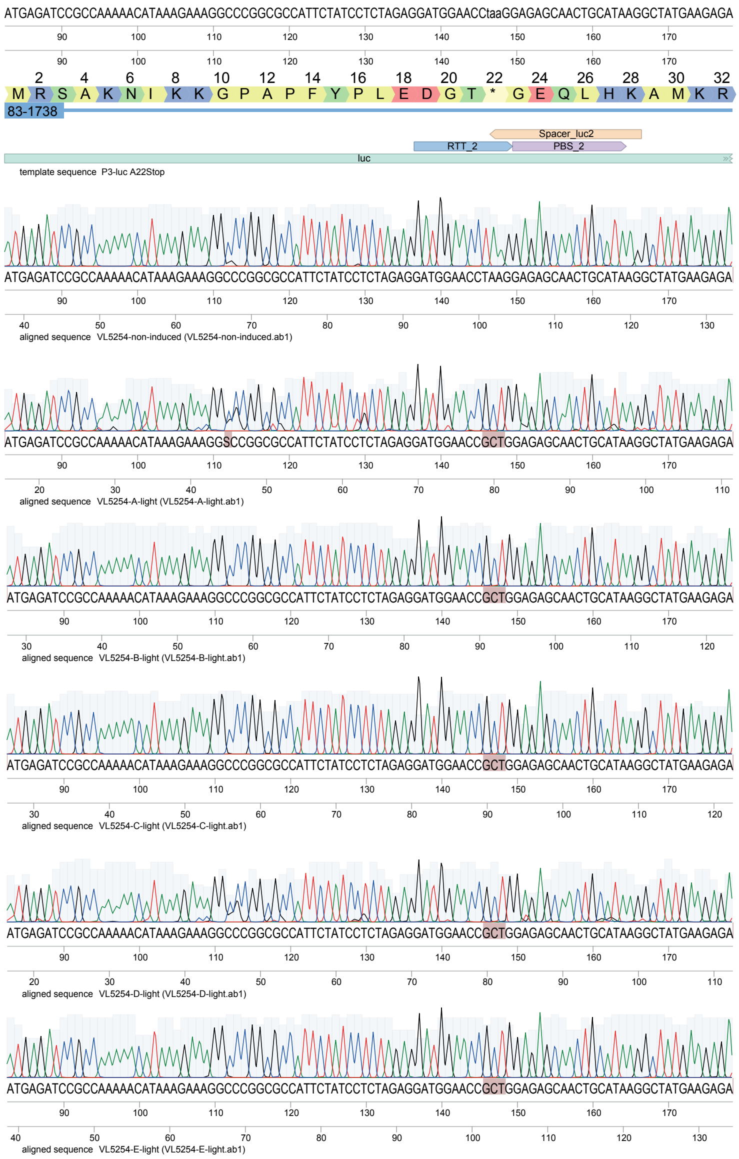

## E. VL5254

template sequence: Luciferase STOP

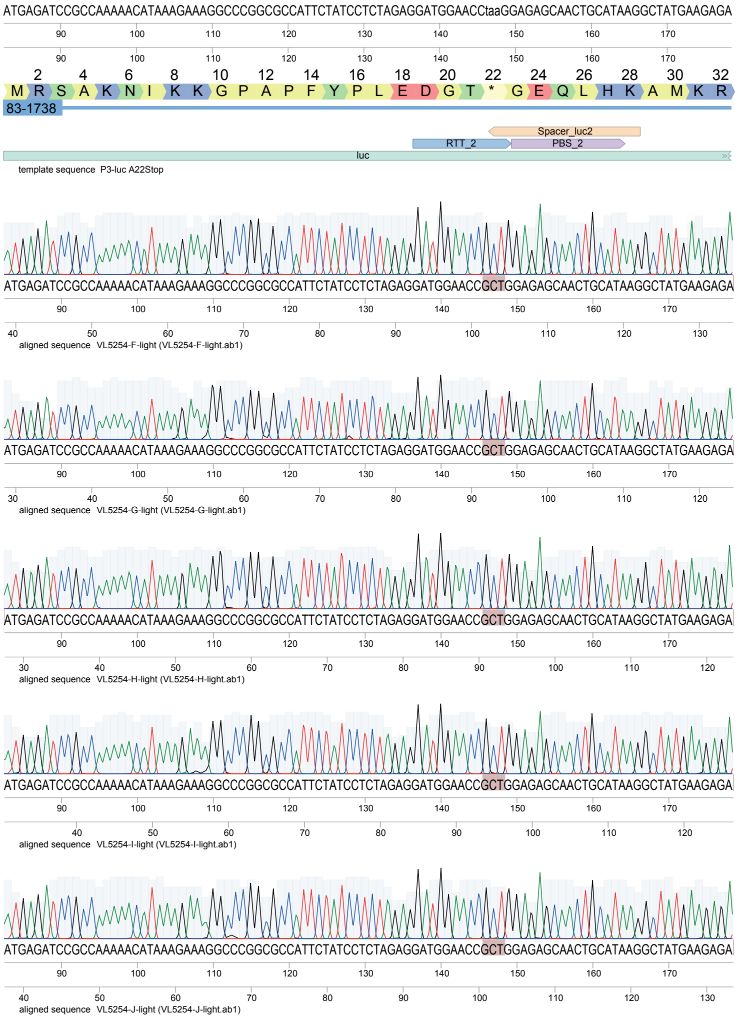

## F. VL5255

template sequence: Luciferase STOP

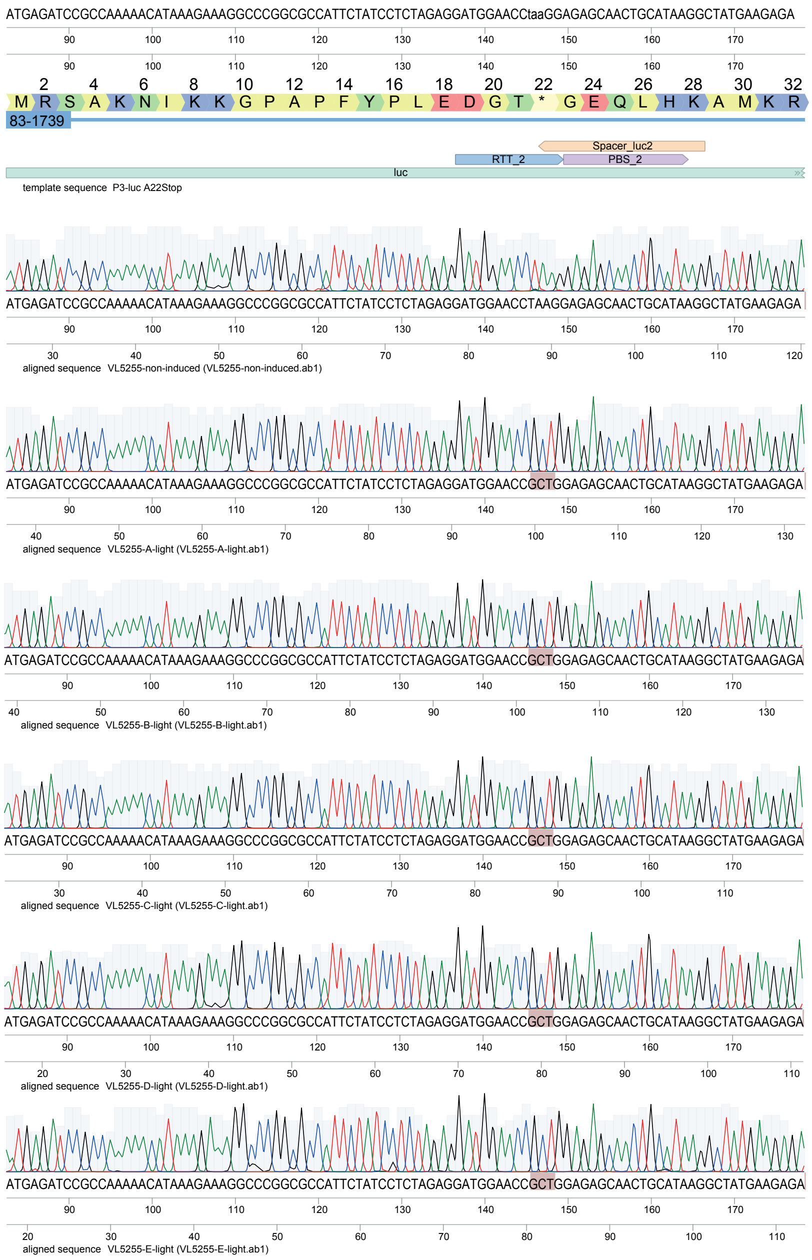

## F. VL5255

template sequence: Luciferase STOP

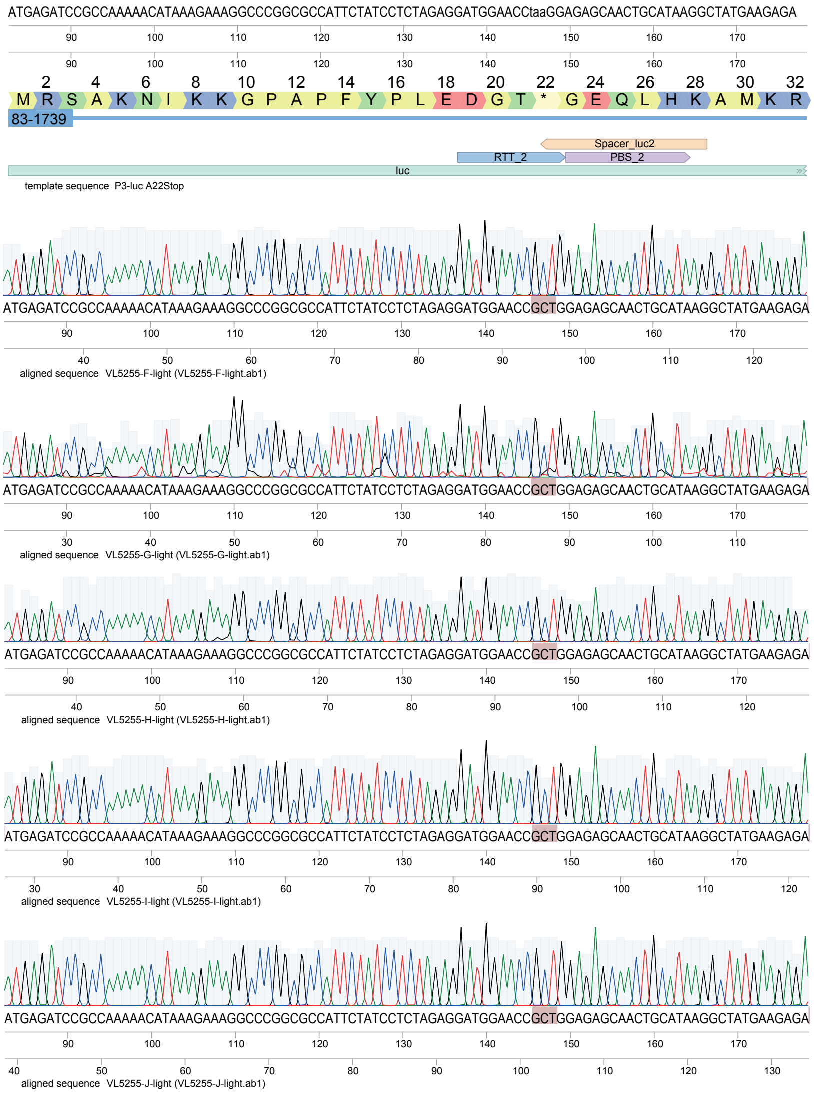

G.

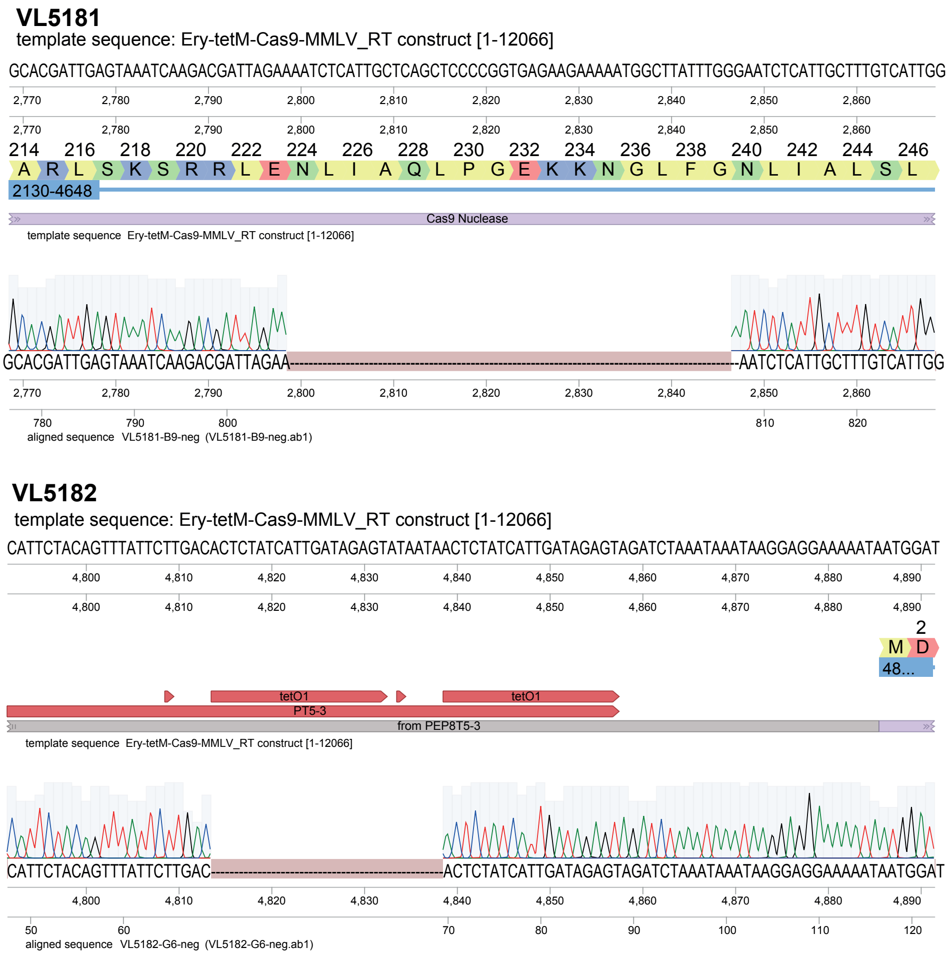


**Supplementary Figure 2.** (**A**) Induction of mbPE2$^{S.pn}$ with aTc, in the presence of targeting pegRNAs, caused severe growth defects (**B**) Comparison of number of colonies (CFU/ml) obtained from plating with or without the inducer (aTc) in presence of targeting pegRNA (VL5182), non-targeting pegRNA (VL7598) or targeting sgRNA (VL7461). **(C-F)** Sanger sequencing of clones from plates containing aTc confirmed that over 85% of colonies had the precise edit as instructed by the pegRNA template. Sanger sequencing of 10 clones is shown for strain VL5182 (C) VL5181 (D) (Luciferase wild type) and VL5254 (E) VL5255 (F) (Luciferase Stop). (**G**) Examples of colonies Sanger sequencing of clones that did survive aTc induction and did not have the desired edit showed the presence of disrupting mutations in the Cas9 domain or Ptet promoter of mbPE2$^{S.pn}$.

#### Supplementary Figure 3.

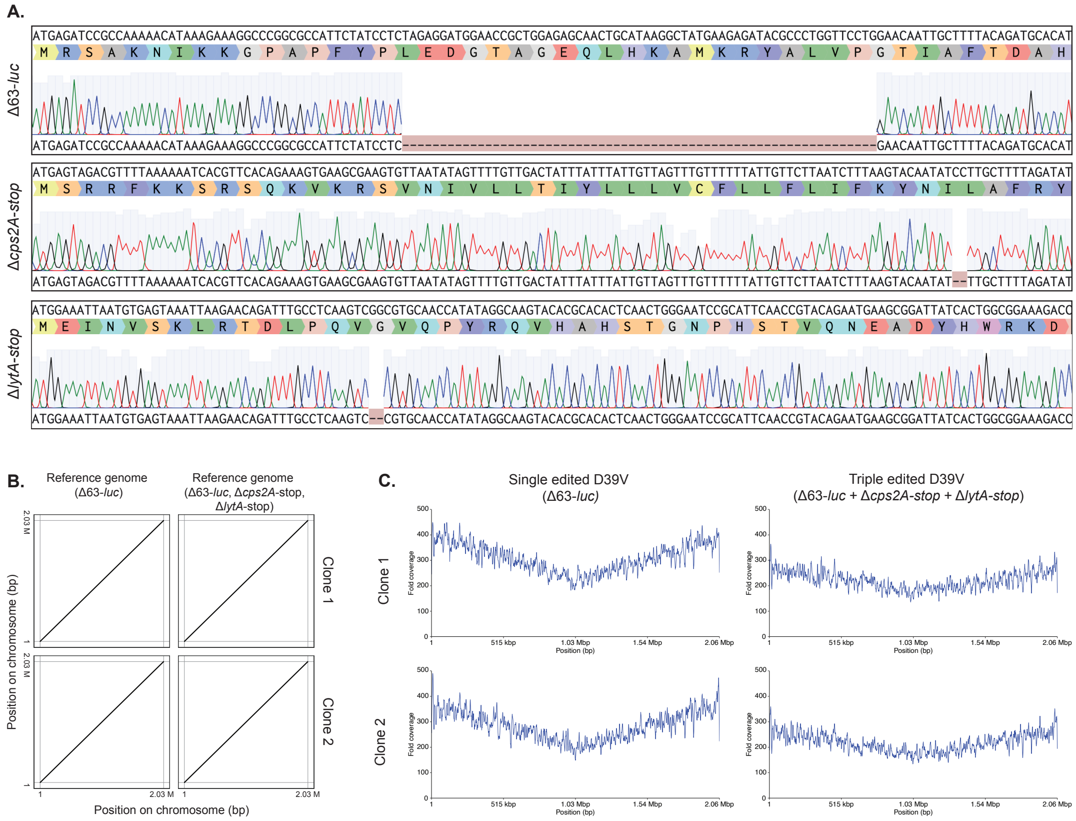


**Supplementary Figure 3. A**) Sanger sequencing chromatograms confirm the deletions in the luciferase, *cps2A*, and *lytA* genes, encoded in the respective pegRNAs. The start codon (ATG) of each gene is positioned at the left hand side, with the translation per triplet shown.
**B)** Alignment of the PacBio assembled genomes of four edited clones to in silico constructed reference sequences showed that no structural rearrangements had occurred in the edited clones.
**C)** Mapping of PacBio reads of four edited clones to in silico constructed reference sequences showed that no duplications had occurred as the result of prime editing.

#### Supplementary Figure 4.

**A.**

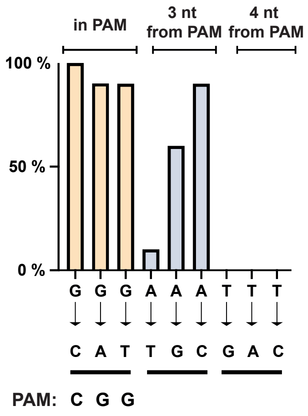


**B.** Template sequence: Substution G -> C

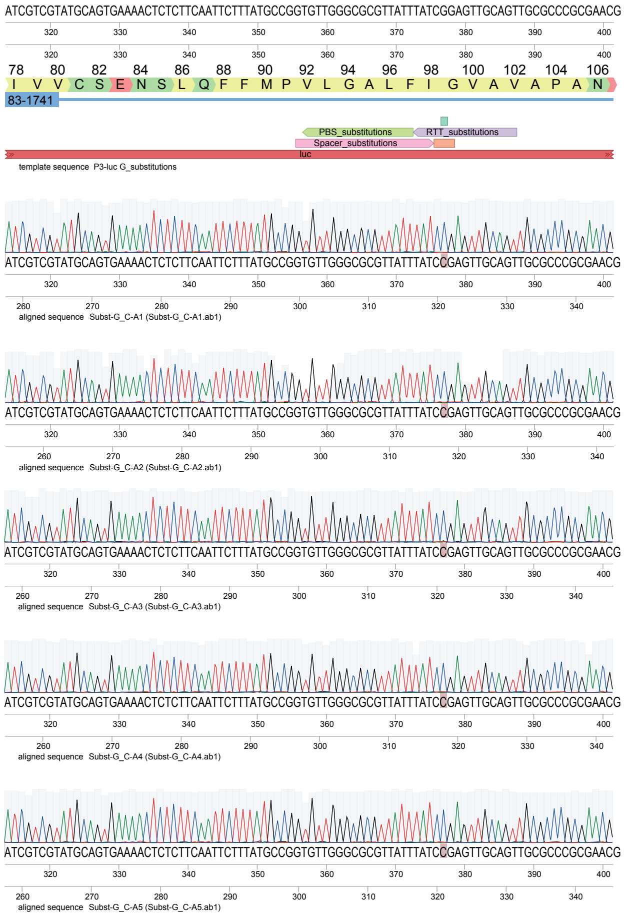

**B.** Template sequence: Substution G -> C

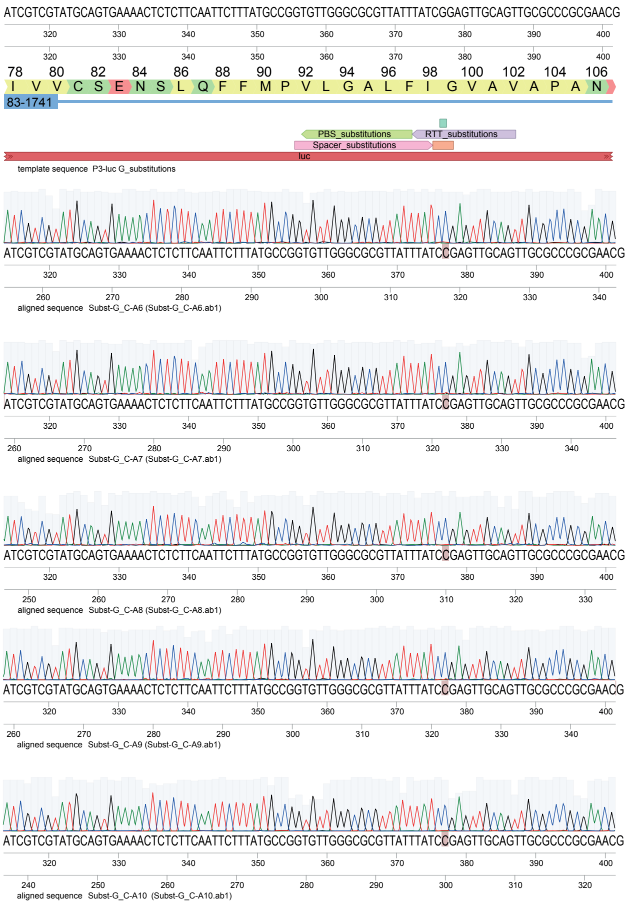


Template sequence: Substution G -> A

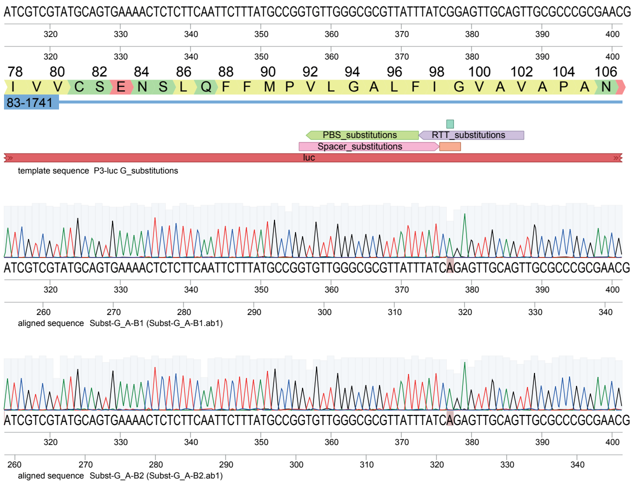

**B.** Template sequence: Substution G -> A

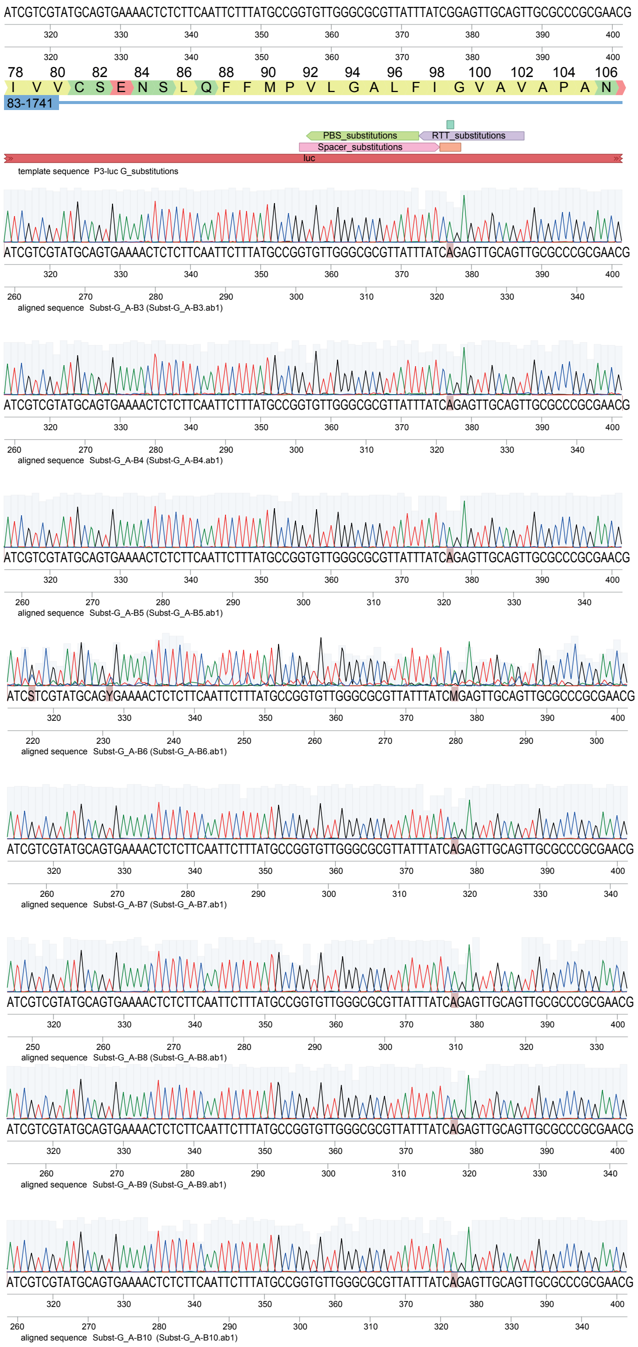

**B.** Template sequence: Substution G -> T

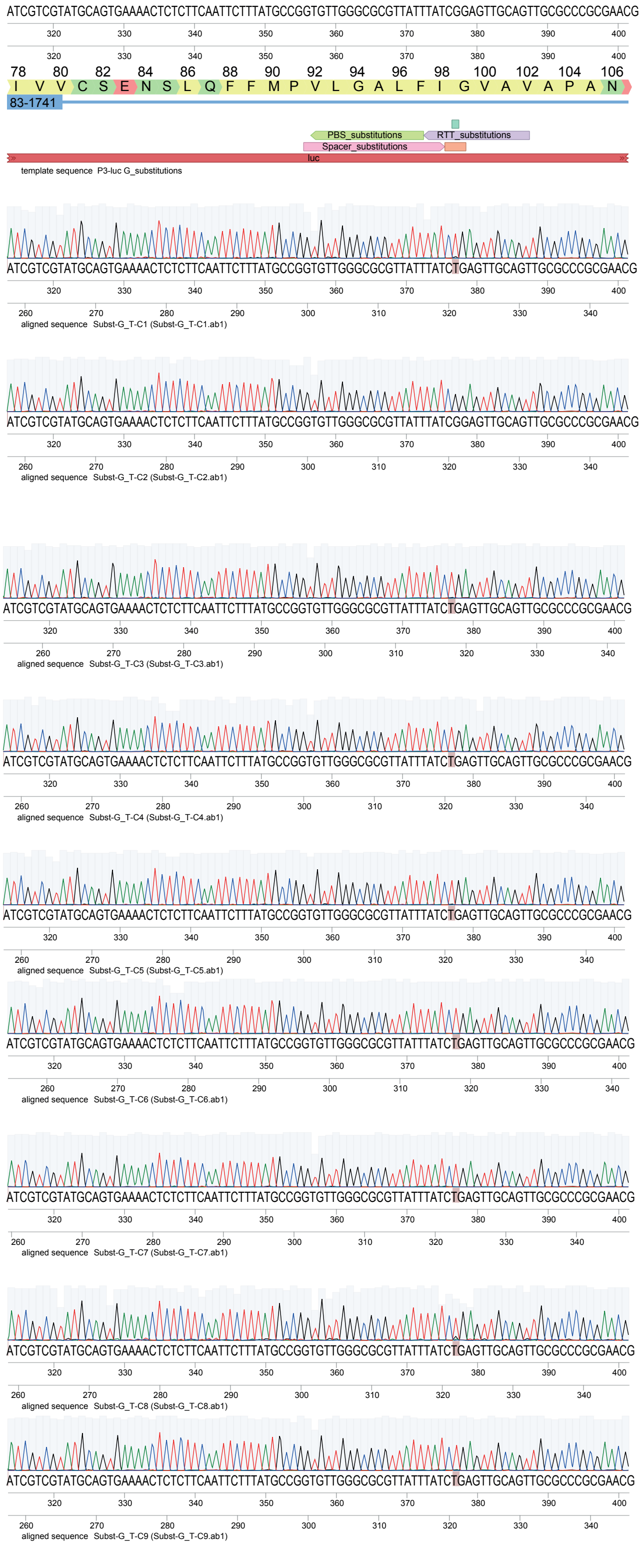

**B.** Template sequence: Substution G -> T

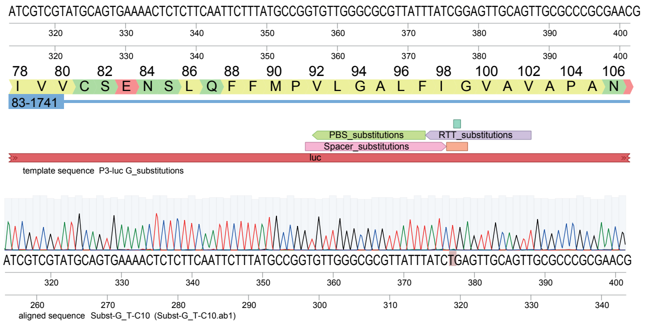


Template sequence: Substution A -> T

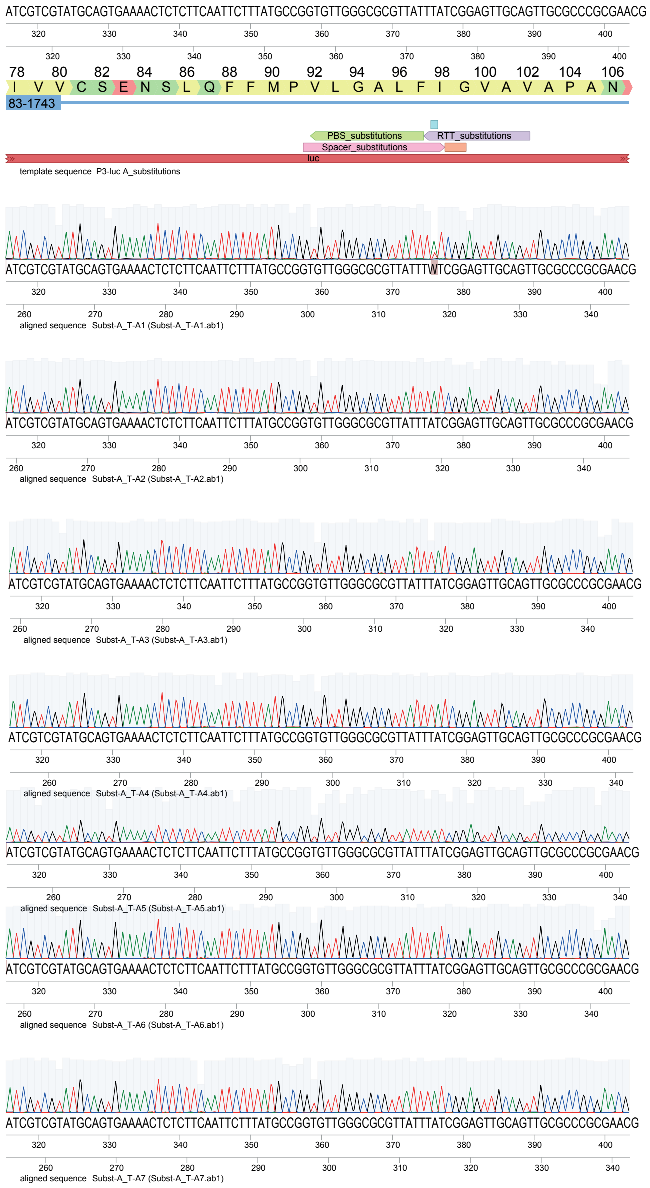

**B.** Template sequence: Substution A -> T

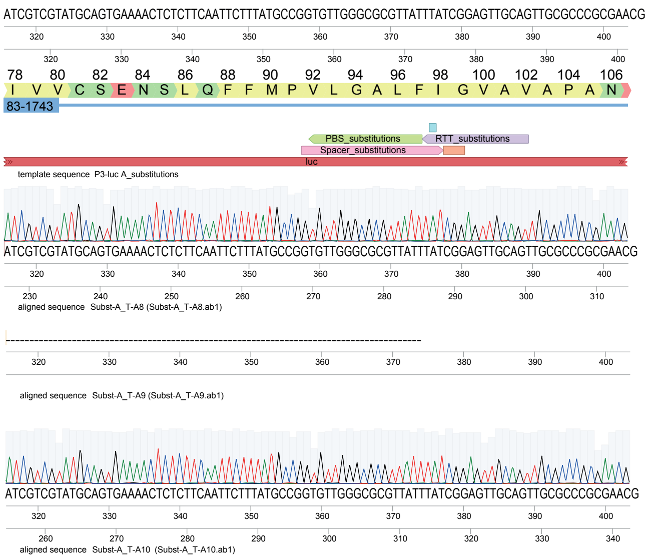


Template sequence: Substution A -> G

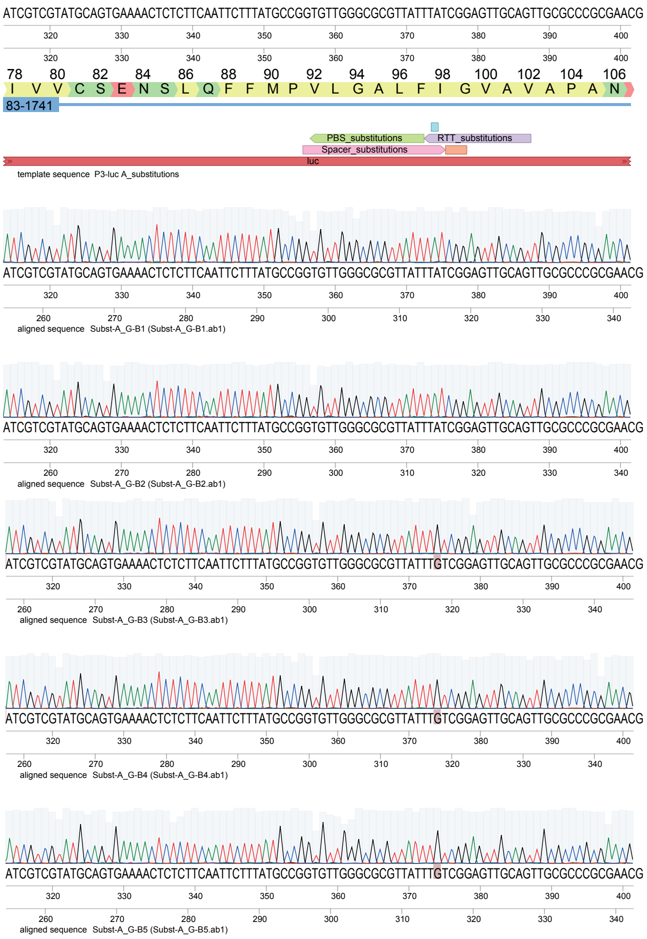

#### B. Template sequence: Substution A -> G

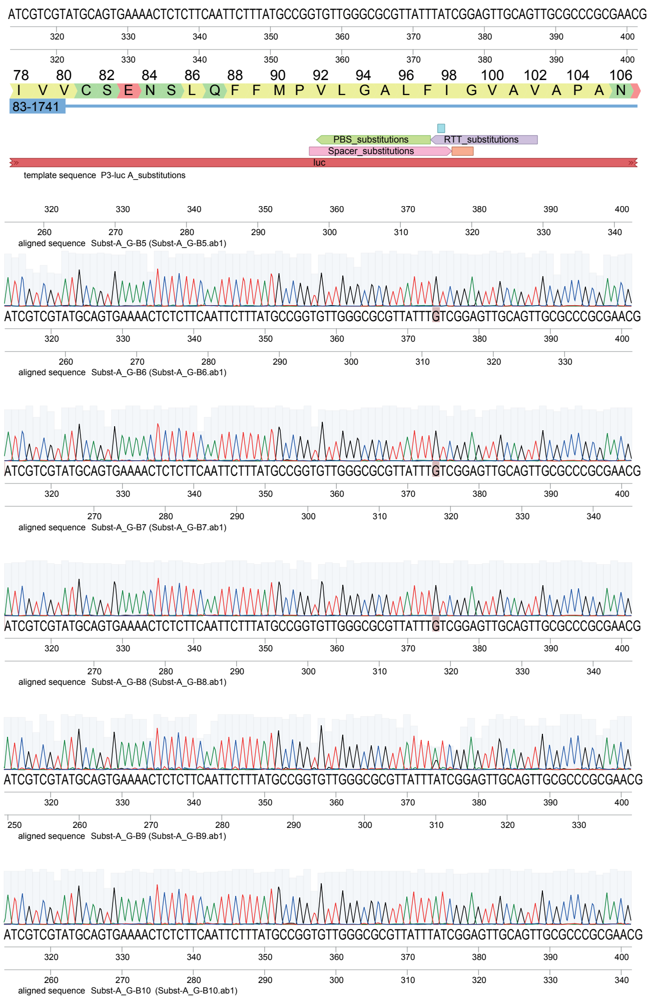


#### B. Template sequence: Substution A -> C

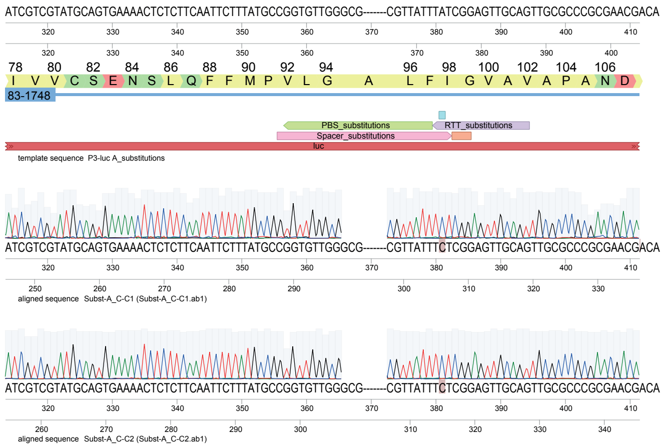

**B.** Template sequence: Substution A -> C

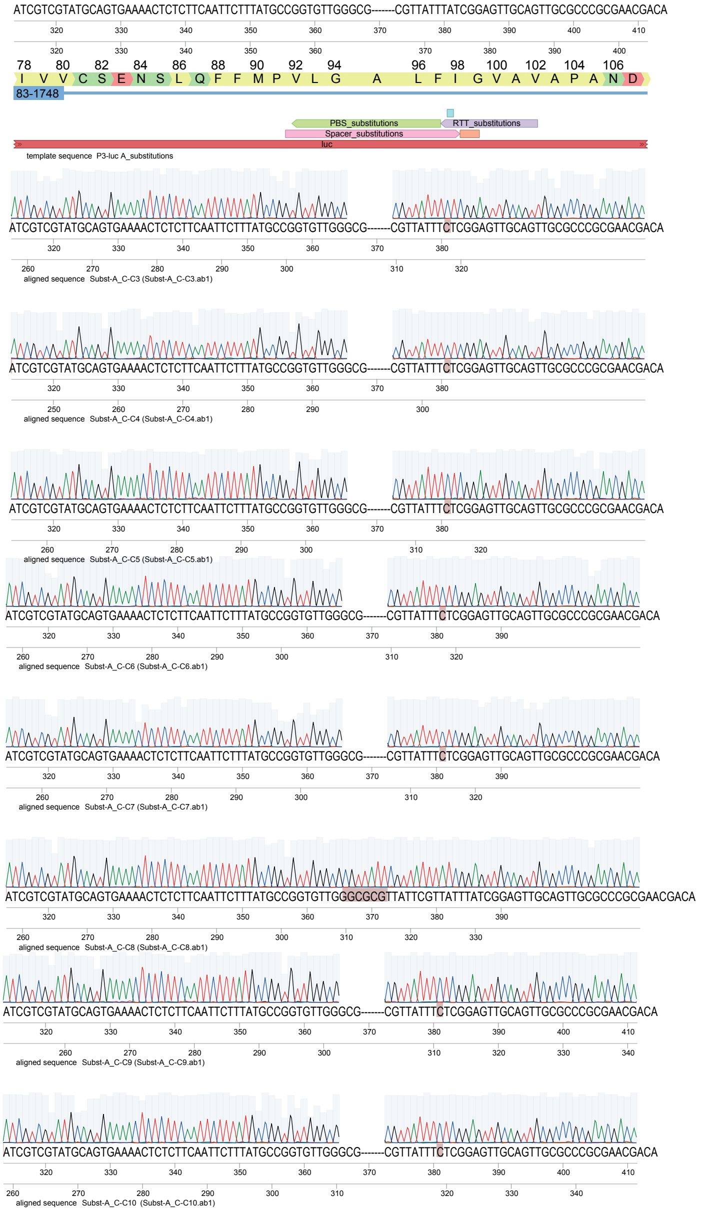

**B.** Template sequence: Substution T -> G

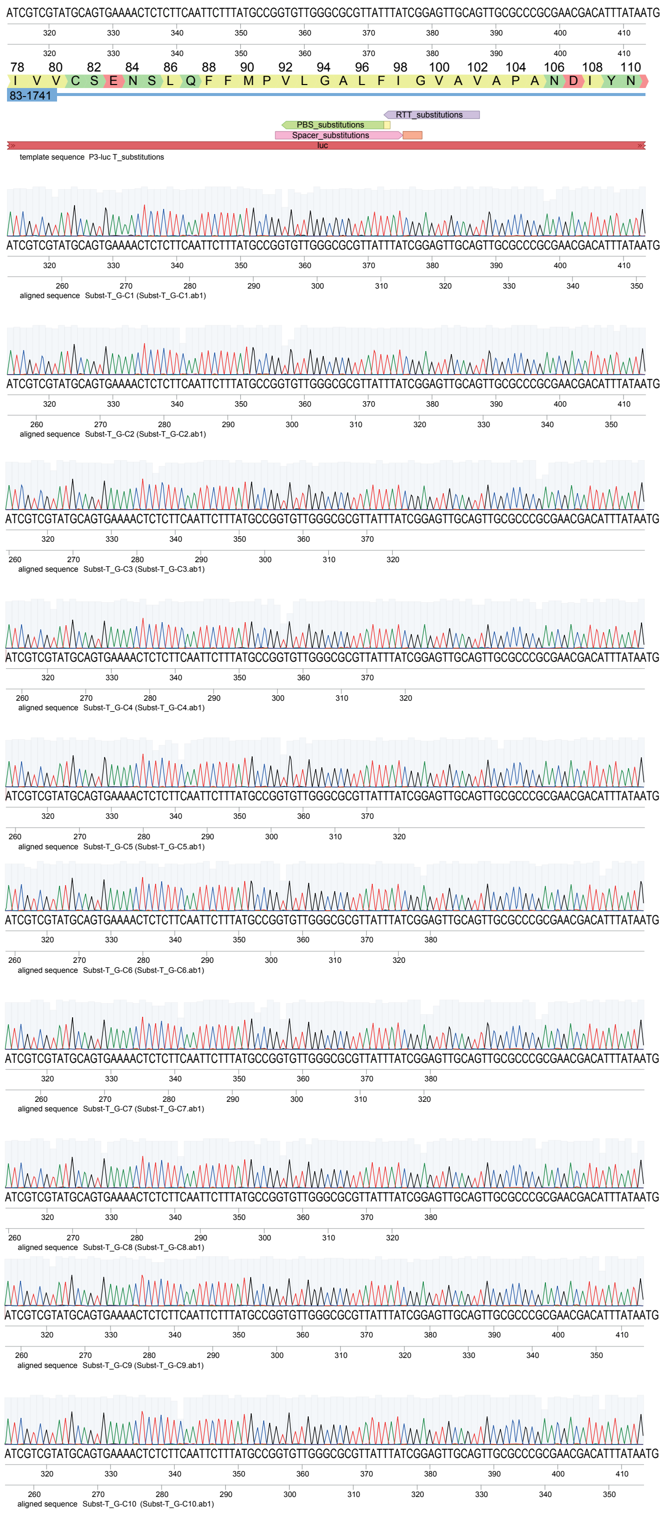

**B.** Template sequence: Substution T -> A

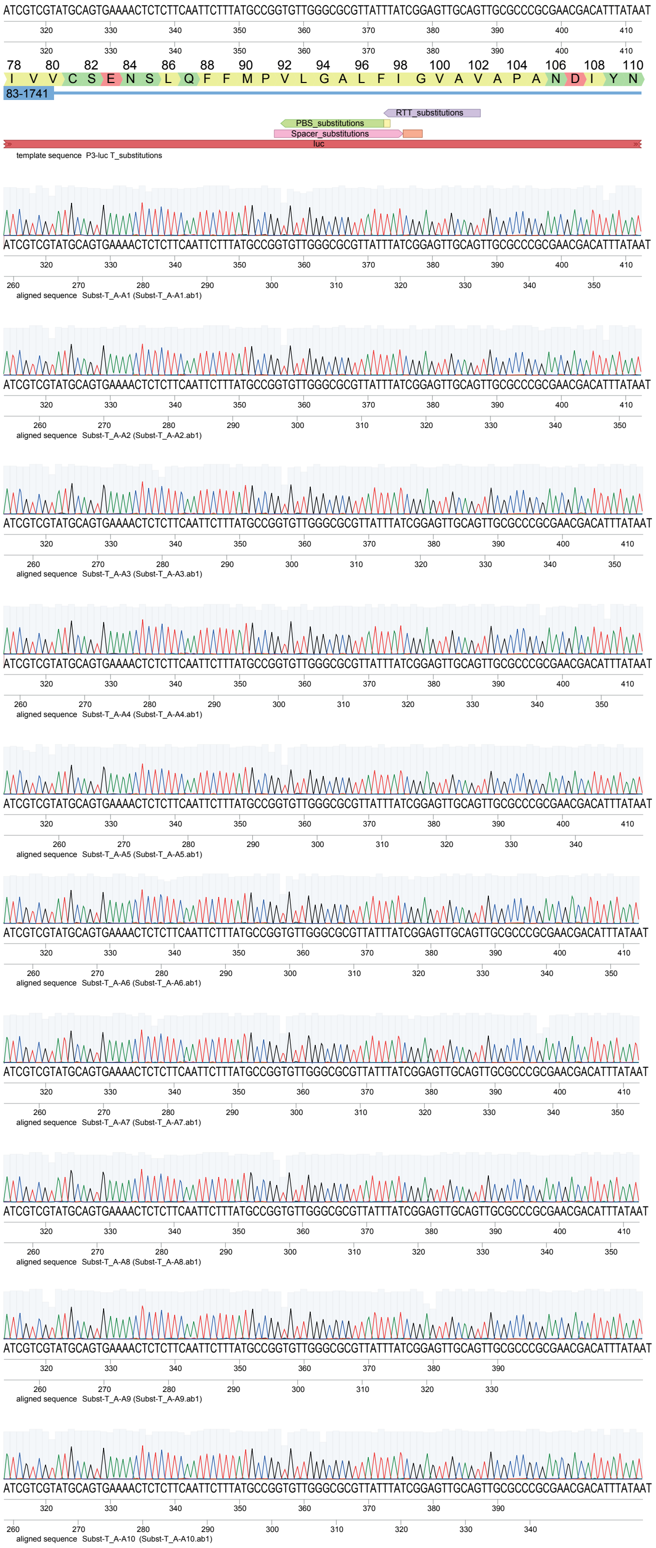

**B.** Template sequence: Substution T -> C

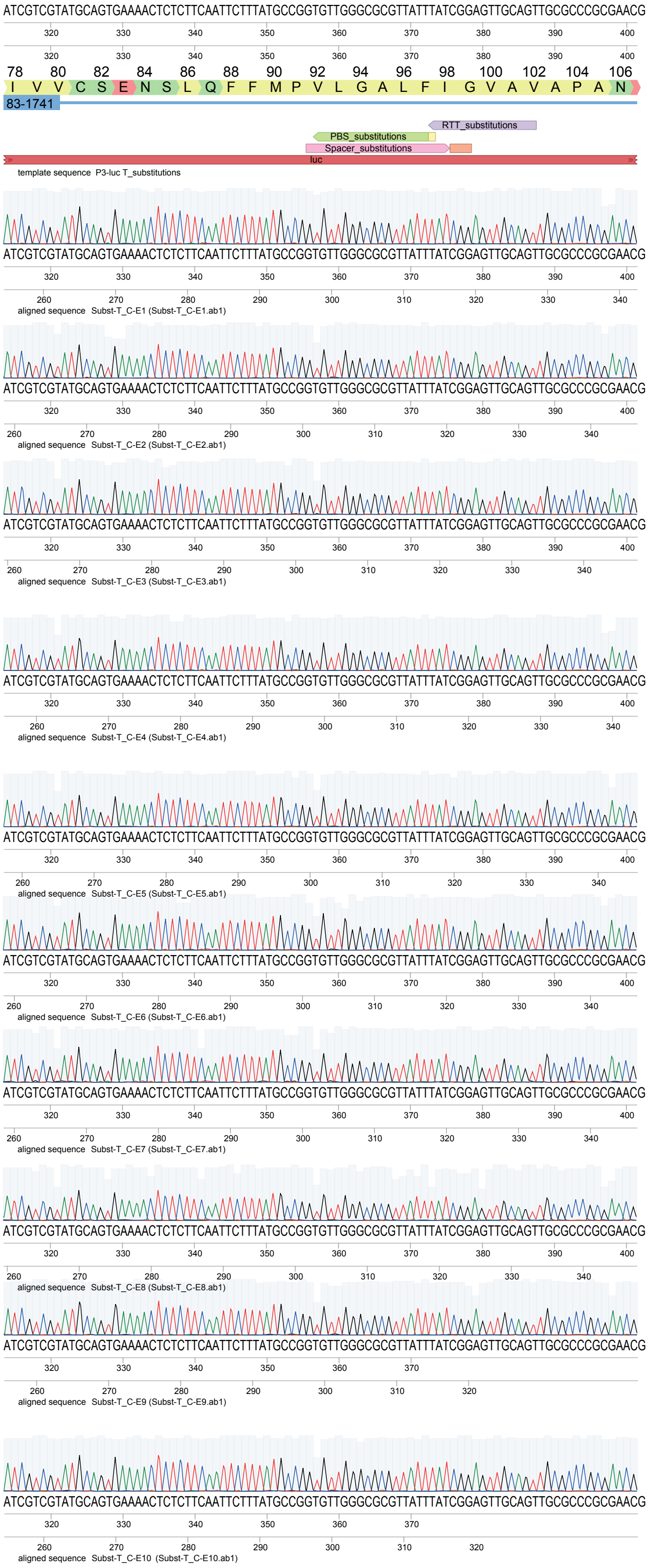

**Supplemental Figure 4.** (**A**) Percentage of successfully edited clones for each substitution of G, T or A disrupting the PAM (G), at 3 (T) and 3 (A) nucleotides distance from the PAM. **(B)** Sanger sequencing profiles of 10 clones for each substitution type used to calculate the percentages in (A).

**Supplementary text**

**Strains construction**

**VL4199**
the strain VL3468 derived from *S. pneumoniae* D39V (lab collection) with a tet-inducible dCas9 (D10A, H840A) integrated at the CEP locus, was used to mutate the alanine residue at position 10 back to an aspartic acid to get a Cas9 nickase (H840A). Two PCR fragments were amplified from this strain: 1) using the forward primer OVL725 included the ISU, tetM, tetR, Ptet and a 31bp fragment of dCas9 with the reverse primer OVL5773 carrying the corresponding nucleotide mutation to get a nickase. 2) the remaining sequence of the Cas9 using the forward primer OVL5774, also carrying the corresponding nucleotide mutation, and the ISD using the primer OVL726. The linker and the MMLVRT sequence were codon-optimized for *S. pneumoniae*. The DNA fragment was obtained as a Gblock (Twist Bioscience) and was amplified using primers OVL1754 and OVL1225. The three DNA fragments were ligated by Golden Gate Assembly (GGA) using AarI. The GGA product was amplified with the flanking primers OVL725 and OVL726, followed by transformation into *S. pneumoniae* D39V. Transformants were selected with tetracycline and confirmed by Sanger sequencing.

**MMLVRT Gblock sequence**
GCAGCCTGGTCTCCGATCGTGTACTGCAGTCTCACCTGCCTGTACTCTGGGGGTTCTTCTG
GAGGATCTTCTGGGTCTGAAACCCCGGGAACGAGCGAGTCAGCGACCCCGGAATCTAGC
GGGGGTTCATCAGGAGGTTCATCTACCCTAAACATCGAGGACGAGTATAGACTACACGA
GACTTCTAAAGAGCCGGACGTGTCACTAGGCAGTACTTGGTTAAGTGATTTCCCTCAGGC
CTGGGCTGAAACAGGGGGGATGGGCCTAGCGGTACGTCAGGCCCCTCTTATCATACCGCT
TAAAGCTACATCAACTCCTGTATCAATTAAACAATACCCAATGTCTCAAGAGGCACGATT
GGGCATTAAGCCGCACATCCAGCGCCTACTAGACCAGGGCATTTTAGTACCATGTCAGTC
TCCATGGAACACCCCGTTACTACCTGTAAAAAAGCCAGGTACTAACGATTATCGTCCAGT
GCAGGACCTAAGAGAGGTTAACAAGCGCGTCGAGGACATACACCCGACTGTACCAAATC
CATACAATTTGTTAAGTGGACTACCTCCTAGTCATCAGTGGTACACTGTACTTGATCTAAA
GGATGCGTTTTTCTGTTTGAGATTGCACCCAACGTCTCAACCATTATTCGCCTTTGAGTGG
AGAGACCCTGAAATGGGCATTTCTGGACAGCTAACGTGGACTCGCCTTCCGCAAGGCTTT
AAAAACAGCCCAACCCTATTCAATGAAGCCCTACATCGCGACTTAGCGGATTTCCGTATC
CAGCATCCGGACTTAATACTACTTCAATACGTGGATGATCTTTTGCTTGCGGCCACCTCAG
AATTGGACTGCCAGCAGGGAACTCGTGCGTTACTACAAACCCTTGGCAACCTAGGGTACC
GAGCAAGCGCAAAGAAGGCGCAGATCTGCCAAAAGCAAGTAAAATACTTAGGTTATCTT
CTTAAGGAGGGGCAACGATGGCTAACGGAAGCCCGAAAGGAAACCGTCATGGGGCAGCC
AACGCCTAAGACACCTCGCCAGTTGCGTGAGTTCCTTGGGAAAGCTGGATTCTGTCGTCT
ATTTATTCCAGGATTTGCTGAGATGGCCGCGCCTCTATATCCACTTACAAAGCCTGGTACC
TTATTTAACTGGGGCCCAGATCAGCAAAAAGCGTATCAAGAGATTAAGCAAGCATTGTTA
ACAGCACCAGCATTAGGGTTACCAGATCTAACTAAACCGTTTGAGCTTTTTGTGGATGAA
AAGCAGGGTTATGCGAAAGGAGTCTTAACCCAGAAATTAGGGCCTTGGCGAAGACCGGT
TGCGTATTTAAGTAAGAAATTGGACCCAGTAGCCGCCGGCTGGCCTCCATGTTTAAGAAT
GGTAGCCGCTATAGCGGTTTTGACTAAGGACGCCGGCAAATTAACTATGGGACAGCCACT
TGTGATCCTTGCGCCGCATGCCGTAGAAGCACTTGTAAAACAGCCGCCAGATCGCTGGCT
ATCTAACGCTAGAATGACGCATTACCAGGCCCTATTATTGGACACAGATCGTGTTCAGTT
CGGGCCGGTTGTAGCATTGAACCCTGCCACACTACTACCTTTACCAGAAGAAGGTCTACA
ACACAACTGCCTTGACATTTTAGCTGAGGCCCATGGCACGCGCCCAGATCTTACCGATCA
GCCGTTGCCGGATGCCGATCACACCTGGTATACGGACGGATCAAGCCTTTTGCAGGAGGG
TCAGCGCAAGGCGGGTGCTGCGGTGACAACGGAAACCGAAGTAATTTGGGCGAAAGCGT
TACCAGCGGGGACCAGCGCTCAACGTGCGGAGTTGATTGCCCTAACGCAAGCACTAAAA
ATGGCTGAAGGAAAGAAGTTGAACGTTTACACGGACAGCCGATATGCTTTCGCGACAGC
TCATATCCACGGCGAAATATATCGTAGAAGAGGCTGGCTAACATCTGAAGGCAAAGAAA
TAAAGAACAAAGACGAGATTTTAGCTTTGCTAAAGGCATTGTTCTTGCCGAAACGTTTGA
GCATCATTCATTGCCCTGGGCACCAGAAGGGTCATTCAGCGGAAGCTCGCGGTAATCGTA
TGGCAGATCAGGCCGCACGCAAAGCGGCTATTACAGAGACTCCAGATACGTCTACGCTAT

TAATTGAGAACTCTTCACCAAGTGGCGGCTCAAAGCGAACAGCCGACGGATCTGAGTTTG
AATAAGGACAGGCAGGTGGCTGAACAAGAGGGCAATCATCGGC

**VL4200**
The GGA assembly product described to obtain the strain VL4199 was used to transform the strain VL956 (*cil::P3-luc , kan*$^R$) from the lab collection. Transformants were selected with tetracycline and confirmed by Sanger sequencing.

**VL4231**
The strain VL4200 was transformed with the plasmid pVL4135. Transformants were selected with spectinomycin and confirmed by Sanger sequencing.

**VL4232**
The strain VL4200 was transformed with the plasmid pVL4134. Transformants were selected with spectinomycin and confirmed by Sanger sequencing.

**VL4297**
To introduce a stop codon in Luc, three fragments were amplified from the strain VL956: 1) ISU, Kanamycin resistance cassette and a fragment of *luc* were amplified with the forward primer OVL3318 and the reverse primer OVL6303, which included the TAA stop codon at residue 22 instead of GCT for alanine. 2) the remaining luc sequence was amplified with the forward oligo OVL6304, also carrying the TAA stop codon, and with the reverse oligo OVL6302. 3) the terminators and the ISD were amplified with the forward primer OVL6080 and the reverse primer OVL3321. The PCR products were ligated by GGA using Esp3I and the product was amplified with the flanking primers OVL 3318 and OVL3321. The resulting PCR product was transformed in the strain VL4199 and the transformants were selected with kanamycin and confirmed by Sanger sequencing.

**VL4347**
The strain VL4297 was transformed with the plasmid pVL4298. Transformants were selected with spectinomycin and confirmed by Sanger sequencing.

**VL4348**
The strain VL4297 was transformed with the plasmid pVL4299. Transformants were selected with spectinomycin and confirmed by Sanger sequencing.

**VL4409**
The strain VL4200 was transformed with the plasmid pVL4403. Transformants were selected with spectinomycin and confirmed by Sanger sequencing.

**VL4449**
The strain VL4297 was transformed with the plasmid pVL4448. Transformants were selected with spectinomycin and confirmed by Sanger sequencing.

**VL4857**
The strain VL1813 from the lab collection (*D39V, hexA::tmp*$^R$) was used to amplify the *hexA::tmp*$^R$ homologous upstream and downstream regions with forward primer OVL7177 and reverse primer OVL7178. The resulting PCR product was transformed in strain VL4231. Transformants were selected with trimethoprim and confirmed by Sanger sequencing.

**VL4858**
The strain VL1813 from the lab collection (*D39V, hexA::tmp*$^R$) was used to amplify the *hexA::tmp*$^R$ homologous upstream and downstream regions with forward primer OVL7177 and reverse primer OVL7178. The resulting PCR product was transformed in strain VL4232. Transformants were selected with trimethoprim and confirmed by Sanger sequencing.

**VL4859**
The strain VL1813 from the lab collection (*D39V, hexA::tmp*$^R$) was used to amplify the *hexA::tmp*$^R$ homologous upstream and downstream regions with forward primer OVL7177 and reverse primer OVL7178. The resulting PCR product was transformed in strain VL4347. Transformants were selected with trimethoprim and confirmed by Sanger sequencing.

**VL4860**
The strain VL1813 from the lab collection (*D39V, hexA::tmp*$^R$) was used to amplify the *hexA::tmp*$^R$ homologous upstream and downstream regions with forward primer OVL7177 and reverse primer OVL7178. The resulting PCR product was transformed in strain VL4348. Transformants were selected with trimethoprim and confirmed by Sanger sequencing.

**VL5149**
To construct the mbPE strain, four fragments were ligated together by GGA using AarI.1) The CEP locus ISU was amplified from strain VL4199 (Sup. table 1) using forward primer OVL7182 and reverse primer OVL7184. 2)The erythromycin resistance cassette was amplified from strain VL3460 from the lab collection (*D39V, Δspv_1662::ery*$^R$) using the forward primer OVL7185 and the reverse primer OV7186. 3) tetM-tetR-$P_{tet}$ and cas9 fragment up to the residue 840 were amplified from strain VL4199 with the forward primer OVL6052 and the reverse primer OVL7187, carrying the CAT codon for histidine instead of GCT for alanine. 4) cas9 fragment from the residue 840 and the linker-MMLVRT sequence were amplified from VL4199 using forward primer OVL6049 carrying the CAT codon for histidine instead of GCT for alanine and reverse primer OVL7183. The GGA product of the four fragments was then amplified with the flanking primers OVL7182 and OVL7183. The resulting PCR product was transformed in D39V. Transformants were selected with erythromycin and confirmed by Sanger sequencing.

**VL5155**
The *cil* locus ISU, *P3-luc* and ISD were amplified from the strain VL4200 using forward primer OVL3318 and reverse primer OVL3321. The resulting PCR product was transformed in the strain VL5149. Transformants were selected with kanamycin and confirmed by Sanger sequencing.

**VL5180**
The *cil* locus ISU, *P3-luc*(A22stop) and ISD were amplified from the strain VL4297 using forward primer OVL3318 and reverse primer OVL3321. The resulting PCR product was transformed in the strain VL5149. Transformants were selected with kanamycin and confirmed by Sanger sequencing.

**VL5181**
The strain VL5155 was transformed with the plasmid pVL4135. Transformants were selected with spectinomycin and confirmed by Sanger sequencing.

**VL5182**
The strain VL5155 was transformed with the plasmid pVL4134. Transformants were selected with spectinomycin and confirmed by Sanger sequencing.

**VL5254**
The strain VL5180 was transformed with the plasmid pVL4298. Transformants were selected with spectinomycin and confirmed by Sanger sequencing.

**VL5255**
The strain VL5180 was transformed with the plasmid pVL4299. Transformants were selected with spectinomycin and confirmed by Sanger sequencing.

**VL5295**
The strain VL5155 was transformed with the plasmid pVL5292. Transformants were selected with spectinomycin and confirmed by Sanger sequencing.

**VL6026**
The strain VL5149 was transformed with the plasmid pVL6005. Transformants were selected with spectinomycin and confirmed by Sanger sequencing.

**VL6125**
To construct VL6125, the mpgA-LgBit-chl$^R$ was cloned in three fragments: 1) mpgA was amplified from D39V with forward primer OVL921 and reverse primer OVL9253. 2) The LgBiT-chl$^R$ sequence was amplified from VL2585 from the lab collection with forward primer OVL9254 and reverse primer OVL9255. 3) the downstream region of mpgA was amplified from D39V with forward primer OVL9256 and reverse primer OVL9257. The three fragments were assembled together by GGA using Esp3I and the resulting product was transformed in the strain VL6220. Transformants were selected with chloramphenicol and confirmed by Sanger sequencing.

**VL6126**
To construct VL6126, the rodZ-LgBit-chl$^R$ was cloned in three fragments: 1) rodZ was amplified from D39V with forward primer OVL9247 and reverse primer OVL9248. 2) The LgBit-chl$^R$ sequence was amplified from VL2585 from the lab collection with forward primer OVL9249 and reverse primer OVL9250. 3) the downstream region of rodZ was amplified from D39V with forward primer OVL9251 and reverse primer OVL5423. The three fragments were assembled together by GGA using Esp3I and the resulting product was transformed in the strain VL6220. Transformants were selected with chloramphenicol and confirmed by Sanger sequencing.

**VL6220**
This strain resulted from the induction of mbPE in the strain VL6026. The correct insertion of SmBit after pbp1a sequence was confirmed by Sanger sequencing.

**VL6247**
To construct VL6247, the hlpA-LgBit-chl$^R$ construct was amplified from the lab collection strain VL2585 with forward primer OVL3895 and reverse primer OVL3900. The resulting product was transformed in the strain VL6220. Transformants were selected with chloramphenicol and confirmed by Sanger sequencing.

**VL6307**
To construct VL6307, the strain VL5155 was transformed with the plasmid pVL6304. Transformants were selected with spectinomycin and confirmed by Sanger sequencing.

**VL6308**
To construct VL6308, the strain VL5155 was transformed with the plasmid pVL6305. Transformants were selected with spectinomycin and confirmed by Sanger sequencing.

**VL6309**
To construct VL6309, the strain VL5155 was transformed with the plasmid pVL6306. Transformants were selected with spectinomycin and confirmed by Sanger sequencing.

**VL6926**
The strain VL6926 results from the mbPE-edition of *luc* in strain VL6307. Clones from mbPE induction experiments were screened by Luciferase activity assay and confirmed by Sanger sequencing.

**VL7244**
To construct VL7244, the strain VL6926 was transformed with the plasmid pVL7230 Transformants were selected with chloramphenicol and confirmed by Sanger sequencing.

**VL7256**

The strain VL7256 results from the mbPE-edition of cps2A (2 bp deletion, leading to a stop codon) in strain VL7244. Clones from mbPE induction experiments were confirmed by Sanger sequencing.

**VL7286**
To construct VL7286, the strain VL7256 was transformed with the plasmid pVL5524. Transformants were selected with spectinomycin and confirmed by Sanger sequencing.

**VL7292**
The strain VL7292 results from the mbPE-edition of lytA (2 bp deletion, leading to a stop codon) in strain VL7286. Clones from mbPE induction experiments were confirmed by Sanger sequencing.

**VL7321**
To construct VL7321, the strain VL5155 was transformed with plasmid pVL7317. Transformants were selected with spectinomycin and confirmed by Sanger sequencing.

**VL7322**
To construct VL7322, the strain VL5155 was transformed with plasmid pVL7318. Transformants were selected with spectinomycin and confirmed by Sanger sequencing.

**VL7323**
To construct VL7323, the strain VL5155 was transformed with plasmid pVL7319. Transformants were selected with spectinomycin and confirmed by Sanger sequencing.

**VL7324**
To construct VL7324, the strain VL5155 was transformed with plasmid pVL7320. Transformants were selected with spectinomycin and confirmed by Sanger sequencing.

**VL7430**
To construct VL7430 the strain VL5155 was transformed with plasmid pVL7370. Transformants were selected with spectinomycin and confirmed by Sanger sequencing.

**VL7431**
To construct VL7431 the strain VL5155 was transformed with plasmid pVL7371. Transformants were selected with spectinomycin and confirmed by Sanger sequencing.

**VL7432**
To construct VL7431 the strain VL5155 was transformed with plasmid pVL7372. Transformants were selected with spectinomycin and confirmed by Sanger sequencing.

**VL7433**
To construct VL7431 the strain VL5155 was transformed with plasmid pVL7373. Transformants were selected with spectinomycin and confirmed by Sanger sequencing.

**VL7434**
To construct VL7431 the strain VL5155 was transformed with plasmid pVL7374. Transformants were selected with spectinomycin and confirmed by Sanger sequencing.

**VL7435**
To construct VL7431 the strain VL5155 was transformed with plasmid pVL7375. Transformants were selected with spectinomycin and confirmed by Sanger sequencing.

**VL7436**
To construct VL7431 the strain VL5155 was transformed with plasmid pVL7376. Transformants were selected with spectinomycin and confirmed by Sanger sequencing.

**VL7437**

To construct VL7431 the strain VL5155 was transformed with plasmid pVL7377. Transformants were selected with spectinomycin and confirmed by Sanger sequencing.

**VL7438**

To construct VL7431 the strain VL5155 was transformed with plasmid pVL7378. Transformants were selected with spectinomycin and confirmed by Sanger sequencing.

**VL7461**

To construct VL7431 the strain VL5155 was transformed with plasmid pVL7456. Transformants were selected with spectinomycin and confirmed by Sanger sequencing.

**VL7487**

To construct VL7487, the recA deletion cassette ΔrecA::tmp was cloned by Golden Gate assembly in three parts. The upstream region of recA was amplified from D39V using primers OVL7147 and OVL11216 and the downstream region using primers OVL11217 and OVL7150. The trimethoprim resistance cassette was amplified from VL4857 using primers OVL11212 and OVL11213. The three fragments were assembled together using SapI and the resulting product was transformed in the strain VL5255. Transformants were selected with trimethoprim and confirmed by Sanger sequencing.

**VL7598**

To construct VL7598, the $P_{tet}$-cas9 sequence from the mbPE construct (VL5255) was cloned by Golden Gate assembly as 2 fragments. The upstream region was amplified using primers OVL5546 and OVL11220 and the downstream region was amplified using primers OVL11221 and OVL726. The two fragments were assembled together using AarI and the resulting product was transformed in D39V. Transformants were selected with erythromycin and confirmed by Sanger sequencing. The resulting strain was then transformed with *cil::P3-luc*(A22stop) as described for the strain VL5180 and with pVL4299 as described for the strain VL5255.

**VL7607**

To construct VL7607 the strain VL5149 was transformed with plasmid pVL4134. Transformants were selected with spectinomycin and confirmed by Sanger sequencing.

**Plasmids construction**

**pVL4132**

To construct pVL4132, the plasmid pVL3991[21] (Addgene plasmid #141090) was amplified with primers OVL5796 and OVL5797. These primers allowed the introduction of Esp3I restriction sites to allow simple cloning of the pegRNAs. The PCR product was assembled by GGA using AarI. The GGA product was transformed in *E. coli* Stbl3 and transformants were selected on LBA plates containing spectinomycin.

**pVL4133**

To construct pVL4133, the plasmid pVL3991[21] (Addgene plasmid #141090) was amplified with primers OVL5798 and OVL5799. These primers allowed the introduction of Esp3I restriction sites to allow simple cloning of the pegRNAs. The PCR product was assembled by GGA using AarI. The GGA product was transformed in *E. coli* Stbl3 and transformants were selected on LBA plates with spectinomycin.

**pVL4134**

To construct pVL4134, the backbone vector pVL4133 was digested with Esp3I. The oligonucleotides for the spacer OVL5891 and OVL5892, the scaffold OVL5893 and OVL5894 and the extension

OVL5895 and OVL5897, were annealed, phosphorylated and ligated with the digested vector. Transformants were selected on LBA plates with spectinomycin.

**pVL4135**

To construct pVL4134, the backbone vector pVL4132 was digested with Esp3I. The oligonucleotides for the spacer OVL5891 and OVL5892, the scaffold OVL5893 and OVL5894 and the extension OVL5895 and OVL5896, were annealed, phosphorylated and ligated with the digested vector. Transformants were selected on LBA plates with spectinomycin.

**pVL4298**

To construct pVL4298, the backbone vector pVL4132 was digested with Esp3I. The oligonucleotides for the spacer OVL6305 and OVL6306, the scaffold OVL5893 and OVL5894 and the extension OVL6307 and OVL6308, were annealed, phosphorylated and ligated with the digested vector. Transformants were selected on LBA plates with spectinomycin.

**pVL4299**

To construct pVL4298, the backbone vector pVL4133 was digested with Esp3I. The oligonucleotides for the spacer OVL6305 and OVL6306, the scaffold OVL5893 and OVL5894 and the extension OVL6307 and OVL6922, were annealed, phosphorylated and ligated with the digested vector. Transformants were selected on LBA plates with spectinomycin.

**pVL4393**

To construct pVL4393, the plasmid pVL4133 was amplified with primers OVL6785 and OVL6788. The PCR product was digested with AarI and ligated to the annealed and phosphorylated oligos for the t-RNA-Asp sequence fragment, OVL6786 and OVL6787. The GGA product was transformed in *E. coli* Stbl3 and transformants were selected on LBA plates with spectinomycin.

**pVL4403**

To construct pVL4403, the backbone vector pVL4393 was digested with Esp3I. The oligonucleotides for the spacer OVL5891 and OVL5892, the scaffold OVL5893 and OVL5894 and the extension OVL5895 and OVL6921, were annealed, phosphorylated and ligated with the digested vector. Transformants were selected on LBA plates with spectinomycin.

**pVL4448**

To construct pVL4448, the backbone vector pVL4393 was digested with Esp3I. The oligonucleotides for the spacer OVL6305 and OVL6306, the scaffold OVL5893 and OVL5894 and the extension OVL6307 and OVL6922, were annealed, phosphorylated and ligated with the digested vector. Transformants were selected on LBA plates with spectinomycin.

**pVL5292**

The cloning of pVL5292 was used as a control for the pegRNA pool cloning strategy. First the vector pVL4134 was amplified with primers OVL7743 and OVL7744, which included the spacer and the scaffold fragments and introduced Esp3I flanking sites. The digested vector was excised and purified from agarose gel. The extension oligonucleotides OVL7747 and OVL7748 were annealed, phosphorylated and ligated with the digested vector. Transformants were selected on LBA plates with spectinomycin.

**pVL5524**

To construct pVL5524, the backbone vector pVL4133 was digested with Esp3I. The oligonucleotides for the spacer OVL8205 and OVL8206, the scaffold OVL5893 and OVL5894 and the extension OVL8207 and OVL8208, were annealed, phosphorylated and ligated with the digested vector. Transformants were selected on LBA plates with spectinomycin.

**pVL5527**

To construct pVL5527, the backbone vector pVL4133 was digested with Esp3I. The oligonucleotides for the spacer OVL8217 and OVL8218, the scaffold OVL5893 and OVL5894 and the extension OVL8219 and OVL8220, were annealed, phosphorylated and ligated with the digested vector. Transformants were selected on LBA plates with spectinomycin.

**pVL5708**
The pPEPY vector (pVL503) was amplified with primers OVL8168 and OVL8169 and the pegRNA for cps2A was amplified from pVL5527 with primers OVL8172 and OVL8179. The PCR products were assembled by GGA using BsaI. Transformants were selected on LBA plates with spectinomycin.

**pVL6005**
To construct pVL6005, the backbone vector pVL4133 was digested with Esp3I. The oligonucleotides for the spacer OVL9026 and OVL9027, the scaffold OVL5893 and OVL5894 and the extension OVL9028 and OVL9029, were annealed, phosphorylated and ligated with the digested vector. Transformants were selected on LBA plates with spectinomycin.

**pVL6304**
To construct pVL6304, first the vector pVL4134 was amplified with primers OVL7743 and OVL7744 which introduced Esp3I flanking sites. The digested vector was excised and purified from agarose gel. The extension oligonucleotides OVL9553 and OVL9554 were annealed, phosphorylated and ligated with the digested vector. Transformants were selected on LBA plates with spectinomycin.

**pVL6305**
To construct pVL6305, first the vector pVL4134 was amplified with primers OVL7743 and OVL7744 which introduced Esp3I flanking sites. The digested vector was excised and purified from agarose gel. The extension oligonucleotides OVL9555 and OVL9556 were annealed, phosphorylated and ligated with the digested vector. Transformants were selected on LBA plates with spectinomycin.

**pVL6306**
To construct pVL6306, first the vector pVL4134 was amplified with primers OVL7743 and OVL7744 which introduced Esp3I flanking sites. The digested vector was excised and purified from agarose gel. The extension oligonucleotides OVL9557 and OVL9558 were annealed, phosphorylated and ligated with the digested vector. Transformants were selected on LBA plates with spectinomycin.

**pVL7227**
To change the resistance marker of pVL4134, the plasmid was amplified with primers OVL10659 and OVL10660. The chloramphenicol resistance cassette was amplified from VL6125 with oligos OVL10661 and OVL10662. The two PCR products were ligated by GGA using AarI. Transformants were selected on LBA plates with chloramphenicol.

**pVL7230**
To construct pVL7230, the backbone vector pVL7227 was digested with Esp3I. The oligonucleotides for the spacer OVL8217 and OVL8218, the scaffold OVL5893 and OVL5894 and the extension OVL8219 and OVL8220, were annealed, phosphorylated and ligated with the digested vector. Transformants were selected on LBA plates with chloramphenicol.

**pVL7317**
To construct pVL7317, the backbone vector pVL4133 was digested with Esp3I. The oligonucleotides for the spacer OVL5891 and OVL5892, the scaffold OVL5893 and OVL5894 and the extension OVL10788 and OVL10789, were annealed, phosphorylated and ligated with the digested vector. Transformants were selected on LBA plates with spectinomycin.

**pVL7318**
To construct pVL7318, the backbone vector pVL4133 was digested with Esp3I. The oligonucleotides for the spacer OVL5891 and OVL5892, the scaffold OVL5893 and OVL5894 and the extension

OVL10790 and OVL10791, were annealed, phosphorylated and ligated with the digested vector. Transformants were selected on LBA plates with spectinomycin.

**pVL7319**
To construct pVL7319, the backbone vector pVL4133 was digested with Esp3I. The oligonucleotides for the spacer OVL5891 and OVL5892, the scaffold OVL5893 and OVL5894 and the extension OVL10792 and OVL10793, were annealed, phosphorylated and ligated with the digested vector. Transformants were selected on LBA plates with spectinomycin.

**pVL7320**
To construct pVL7320, the backbone vector pVL4133 was digested with Esp3I. The oligonucleotides for the spacer OVL5891 and OVL5892, the scaffold OVL5893 and OVL5894 and the extension OVL10794 and OVL10795, were annealed, phosphorylated and ligated with the digested vector. Transformants were selected on LBA plates with spectinomycin.

**pVL7370**
To construct pVL7370, the backbone vector pVL4133 was digested with Esp3I. The oligonucleotides for the spacer OVL11019 and OVL11020, the scaffold OVL5893 and OVL5894 and the extension OVL11021 and OVL11022, were annealed, phosphorylated and ligated with the digested vector. Transformants were selected on LBA plates with spectinomycin.

**pVL7371**
To construct pVL7370, the backbone vector pVL4133 was digested with Esp3I. The oligonucleotides for the spacer OVL11019 and OVL11020, the scaffold OVL5893 and OVL5894 and the extension OVL11023 and OVL11024, were annealed, phosphorylated and ligated with the digested vector. Transformants were selected on LBA plates with spectinomycin.

**pVL7372**
To construct pVL7370, the backbone vector pVL4133 was digested with Esp3I. The oligonucleotides for the spacer OVL11019 and OVL11020, the scaffold OVL5893 and OVL5894 and the extension OVL11025 and OVL11026, were annealed, phosphorylated and ligated with the digested vector. Transformants were selected on LBA plates with spectinomycin.

**pVL7373**
To construct pVL7370, the backbone vector pVL4133 was digested with Esp3I. The oligonucleotides for the spacer OVL11019 and OVL11020, the scaffold OVL5893 and OVL5894 and the extension OVL11027 and OVL11028, were annealed, phosphorylated and ligated with the digested vector. Transformants were selected on LBA plates with spectinomycin.

**pVL7374**
To construct pVL7370, the backbone vector pVL4133 was digested with Esp3I. The oligonucleotides for the spacer OVL11019 and OVL11020, the scaffold OVL5893 and OVL5894 and the extension OVL11029 and OVL11030, were annealed, phosphorylated and ligated with the digested vector. Transformants were selected on LBA plates with spectinomycin.

**pVL7375**
To construct pVL7370, the backbone vector pVL4133 was digested with Esp3I. The oligonucleotides for the spacer OVL11019 and OVL11020, the scaffold OVL5893 and OVL5894 and the extension OVL11031 and OVL11032, were annealed, phosphorylated and ligated with the digested vector. Transformants were selected on LBA plates with spectinomycin.

**pVL7376**
To construct pVL7370, the backbone vector pVL4133 was digested with Esp3I. The oligonucleotides for the spacer OVL11019 and OVL11020, the scaffold OVL5893 and OVL5894 and the extension

OVL11033 and OVL11034, were annealed, phosphorylated and ligated with the digested vector. Transformants were selected on LBA plates with spectinomycin.

**pVL7377**
To construct pVL7370, the backbone vector pVL4133 was digested with Esp3I. The oligonucleotides for the spacer OVL11019 and OVL11020, the scaffold OVL5893 and OVL5894 and the extension OVL11035 and OVL11036, were annealed, phosphorylated and ligated with the digested vector. Transformants were selected on LBA plates with spectinomycin.

**pVL7378**
To construct pVL7370, the backbone vector pVL4133 was digested with Esp3I. The oligonucleotides for the spacer OVL11019 and OVL11020, the scaffold OVL5893 and OVL5894 and the extension OVL11037 and OVL11038, were annealed, phosphorylated and ligated with the digested vector. Transformants were selected on LBA plates with spectinomycin.

**pVL7456**
To construct pVL7456, pVL3991 (Addgene plasmid #141090) was digested using EcoRV, and the backbone amplified using primers OVL6185 and OVL6186. Oligonucleotides OVL1020 and OVL1021, encoding the sgRNA targeting firefly luciferase, were annealed in TEN buffer, and ligated into the amplified pVL3991 backbone through a Golden Gate reaction using Esp3I. Transformants were select on LBA plates with spectinomycin.
